# Molecular Dynamics Activation of γ-Secretase for Cleavage of Notch1 Substrate

**DOI:** 10.1101/2023.09.26.559539

**Authors:** Hung N. Do, Shweta R. Malvankar, Michael S. Wolfe, Yinglong Miao

## Abstract

γ-Secretase is an intramembrane aspartyl protease complex which cleaves the transmembrane domain of over 150 peptide substrates, including amyloid precursor protein (APP) and the Notch family of receptors, via two conserved aspartates D257 and D385 in the Presenilin-1 (PS1) catalytic subunit. However, while the activation of γ-secretase for cleavage of APP has been widely studied, the cleavage of Notch by γ-secretase remains poorly explored. Here, we combined Gaussian accelerated Molecular Dynamics (GaMD) simulations and mass spectrometry (MS) analysis of proteolytic products to present the first dynamic models for cleavage of Notch by γ-secretase. MS showed that γ-secretase cleaved the WT Notch at Notch residue G34, while cleavage of L36F mutant Notch occurred at Notch residue C33. Initially, we prepared our simulation systems starting from the cryoEM structure of Notch-bound γ-secretase (PDB: 6IDF) and failed to capture the proper cleavages of WT and L36F Notch by γ-secretase. We then discovered an incorrect registry of the Notch substrate in the PS1 active through alignment of the experimental structure of Notch-bound (PDB: 6IDF) and APP-bound γ-secretase (PDB: 6IYC). Every residue of APP substrate was systematically mutated to the corresponding Notch residue to prepare a resolved model of Notch-bound γ-secretase complexes. GaMD simulations of the resolved model successfully captured γ-secretase activation for proper cleavages of both WT and L36F mutant Notch. Our findings here provided mechanistic insights into the structural dynamics and enzyme-substrate interactions required for γ-secretase activation for cleavage of Notch and other substrates.

## Introduction

γ-Secretase, “the proteasome of the membrane”^1^, is an intramembrane aspartyl protease complex composed of four components Nicastrin (NCT), Aph-1, Pen-2, and Presenilin-1 (PS1)^2,3^. γ-Secretase carries out its proteolytic activity via two conserved aspartates (D257 and D385) in the PS1 catalytic subunits^4, 5^, cleaving more than 150 peptide substrates^6^, including amyloid precursor protein (APP) and the Notch family of cell-surface receptors. However, while the cleavage of APP by γ-secretase has been widely studied^7-9^, the proteolysis of other substrates, including Notch, by γ-secretase remains underexplored.

Molecular dynamics (MD) is a powerful computational technique for simulating biomolecular dynamics at an atomistic level^10^. However, conventional MD (cMD) is often limited to microsecond simulation timescale, and thus cannot sufficiently sample many biological processes of interest^11^. Enhanced sampling techniques have been developed to overcome this challenge. Gaussian accelerated molecular dynamics (GaMD) is an enhanced sampling technique that works by applying a harmonic boost potential to smooth biomolecular potential energy surface^12^. Since this boost potential exhibits a near Gaussian distribution, cumulant expansion to the second order (“Gaussian approximation”) can be applied to achieve proper energetic reweighting^13^. GaMD allows for simultaneous unconstrained enhanced sampling and free energy calculations of large biomolecules^12^. GaMD^12^ and its variant methods^14-19^ have been successfully demonstrated on enhanced sampling of ligand binding, protein/RNA folding, protein conformational, as well as protein-membrane, protein-protein, and protein-nucleic acid interactions^11^.

In 2020, our labs^8^ combined complementary GaMD simulations and biochemical experiments to investigate mechanisms of the γ-secretase activation and the initial endoproteolytic (ε) cleavage of wildtype (WT) and familial Alzheimer’s disease (FAD)-mutant forms of APP substrate. GaMD simulations captured spontaneous activation of γ-secretase: First, the protonated D257 formed a hydrogen bond with the backbone carboxyl group of APP residue L49. Then, a water molecule was recruited between the two catalytic aspartates through hydrogen bonds. In this way, the water molecule was activated for nucleophilic attack on the carbonyl carbon of APP residue L49 to carry out ε cleavage. GaMD simulations also revealed that APP FAD mutations I45F and T48P preferred ε cleavage at the L49–V50 amide bond, whereas M51F shifted the ε cleavage site to the T48–L49 amide bond, results highly consistent with experimental analyses of APP proteolytic products using mass spectrometry (MS) and western blotting^2, 8^. We also combined Peptide GaMD (Pep-GaMD) simulations with MS and western blotting to investigate tripeptide trimming of wildtype (WT) and FAD-mutant Aβ49 substrates by γ-secretase^20^. The Pep-GaMD simulations revealed remarkable structural rearrangements of both γ-secretase and Aβ49, where hydrogen-bonded catalytic aspartates and water were poised to carry out the ζ cleavage of Aβ49 to Aβ46. Furthermore, this trimming step required inclusion of ε coproduct APP intracellular domain (AICD) with a positively charged N-terminus. These simulation findings were also highly consistent with biochemical experimental data^20, 21^. Very recently, we performed complimentary GaMD simulations alongside biochemical experiments to determine the effects of PS1 FAD mutations on γ-secretase activation for ε cleavage of APP^9^. In that study, we captured a similar mechanism of WT γ-secretase activation for ε cleavage of APP as Bhattarai et al.^8^, even though a different catalytic aspartate was protonated (D385 compared to D257)^9^.

In this work, we combined MS analyses and GaMD simulations to capture the activation of γ-secretase for cleavage of WT and L36F Notch. MS showed that γ-secretase cleaves WT Notch at Notch residue G34 and L36F mutant Notch at Notch residue C33. We failed to capture the experimentally determined cleavages of WT and L36F Notch by γ-secretase in the GaMD simulations starting from the cryoEM structure of Notch-bound γ-secretase (PDB: 6IDF^22^). We then discovered an incorrect registry of the Notch substrate in the PS1 active site by structurally aligning the PDB structures of Notch-bound (PDB: 6IDF^22^) and APP-bound γ-secretase (PDB: 6IYC^23^). Eventually, we mutated every residue of APP substrate to the corresponding Notch residue to prepare a resolved model Notch-bound γ-secretase complexes and successfully captured the correct cleavages of WT and L36F Notch at Notch residue G34 and C33, respectively, by γ-secretase in our GaMD simulations. Our GaMD simulations and MS experiments provided mechanistic insights into the structural dynamics and enzyme-substrate interactions required for γ-secretase activation for cleavage of Notch and other substrates.

## Materials and Methods

### Generation of WT and L36F constructs

A truncated WT Notch substrate construct was designed with a 6XHis epitope tag on N-terminus and FLAG tag on the C-terminus in the pET vector (gift of C. Sanders, Vanderbilt University). The WT Notch construct was then mutated using designed primers and Quik Change Lightning mutagenesis kit (Agilent) according to the manufacturer’s instruction to give Notch_L36F construct, and the mutation was verified by sequencing (ACGT, Inc).

### Purification of WT and L36F Notch

*E. coli* BL21 cells were transformed with the respective Notch constructs and plated on LB-ampicillin agar plates and incubated at 37 °C overnight. A single colony was picked for each construct variant and grown with shaking in LB media at 37 °C until OD600 reached 0.8. Cells were induced with 0.5 mM IPTG and were grown for 3 h. Cells were then collected by centrifugation and resuspended in 50 mM HEPES pH 8, 1% Triton X-100. The cell suspension was passed through a French press, and the lysate was incubated with anti-FLAG M2-agarose beads from Sigma-Aldrich. Bound substrates were then eluted from the beads with 100 mM glycine buffer, pH 2.7, with 0.25% NP-40 detergent and then stored at −80 °C. The purified proteins were confirmed by MALDI-TOF MS and SDS-PAGE.

### WT γ-secretase expression and purification

γ-Secretase was expressed in HEK 293 F cells by transfection with tetracistronic pMLINK vector encoding all four components of the protease complex (gift of Y. Shi, Tsinghua University)^24^. For transfection, HEK 293 F cells were grown in suspension in unsupplemented Freestyle 293 media (Life Technologies, 12338-018) until cell density reached 2 × 10^6^ cells/mL. Tetracistronic pMLINK vector (150 mg) was mixed with 25 kDa linear polyethylemimines (PEI; 450 mg) and incubated for 30 min at room temperature. The DNA-PEI mixtures were then added to HEK cells, and cells were grown for 60 h. Cells were harvested, and γ-secretase was purified as described previously^25^.

### Reaction of γ-Secretase and Notch Substrate

The purified γ-secretase containing presenilin-1 was incubated for 30 min at 37 °C in the assay buffer containing 50 mM 4-(2-hydroxyethyl)-1-piperazineethanesulfonic acid (HEPES, pH 7.0), 150 mM NaCl, 0.1% 1,2-dioleoyl-sn-glycero-3-phosphocholine (DOPC), 0.025% 1,2-dioleoyl-sn-glycero-3-phosphoethanolamine (DOPE), and 0.25% zwitterionic detergent 3-[(3-cholamidopropyl)dimethylammonio]-2-hydroxy-1-propanesulfonate (CHAPSO), at final enzyme concentration 30 nM. The Notch substrates were then added to final concentrations of 20 uM in the respective reactions, and the reaction mixtures were incubated at 37 °C for 4h.

### Detection of NICD-like products

The truncated NICD-like proteolytic fragment, containing a C-terminal FLAG epitope tag, produced from γ-secretaseenzymatic reactions with WT or L36F Notch-based substrate was isolated by immunoprecipitation with anti-FLAG M2 beads (Sigma-Aldrich) in 10 mM MES (2-(4-morpholino) ethanesulfonic acid) pH 6.5, 10 mM NaCl, 0.05% DDM detergent for 16 hours at 4 °C. NICD-like products were eluted from the anti-FLAG beads with acetonitrile: water (1:1) with 0.1% trifluoroacetic acid (TFA). The elutes were run on a Bruker Autoflex maX MALDI-TOF mass spectrometer.

### Simulation system setup

First, we used the cryoEM structure of Notch-bound γ-secretase (PDB: 6IDF)^22^ to prepare the cryoEM simulation systems. The artificial disulfide bonds between C112 of PS1-Q112C and C9 of Notch-P9C were removed as the WT residues (Q112 and P9) were restored. Second, we built the model WT Notch-bound γ-secretase by computationally mutating every APP residue in APP-bound γ-secretase (PDB: 6IYC)^23^ to Notch residue using the *Mutation* function of CHARMM-GUI^26-32^ as following:

APP LVFFAEDVGSNKGAIIGLMVGGVVIATVIVITLVMLKKK

Notch VQSETVEPPPPAQLHFMYVAAAAFVLLFFVGCGVLLSRK

The large hydrophilic loop that connected TM6a and TM7 was not modelled based on previous studies^8, 9, 20^. The L36F Notch mutations for both cryoEM and model Notch-bound γ-secretase were computationally generated using the *Mutation* function of CHARMM-GUI^26-32^. We chose to protonate residue D385 in PS1 to simulate the activation of γ-secretase for cleavage of Notch based on our previous study^9^. All heteroatom molecules were removed, and all chain termini were capped with neutral patched (acetyl and methylamide). The enzyme-substrate complexes were embedded in POPC membrane lipid bilayers and then solvated in 0.15 M NaCl solutions using the CHARMM-GUI webserver^26-32^.

### Simulation protocols

The CHARMM36m force field parameter set^33^ was used for the protein and lipids. Periodic boundary conditions were applied for the simulation systems. Bonds containing hydrogen atoms were restrained with the SHAKE^34^ algorithm, and a 2-fs time step was used. The temperature was kept at 310 K using the Langevin thermostat^35, 36^ with a friction coefficient of 1.0 ps^-1^. The pressure was kept constant at 1.0 bar using the Berendsen barostat^37^ with semi-isotropic coupling with enabled surface tension in the X-Y plane. The Berendsen coupling constant was set to 0.5 ps. The electrostatic interactions were calculated using the particle mesh Ewald (PME) summation^38^ with a cutoff of 9.0 Å for long-range interactions. The simulation systems were initially energetically minimized for 5000 steps using the steepest-descent algorithm and equilibrated with the constant number, volume, and temperature (NVT) ensemble at 310 K. They were further equilibrated for 375 ps at 310 K with the constant number, pressure, and temperature (NPT) ensemble. Short cMD simulations were then performed for 10 ns using the NPT ensemble with constant surface tension at 1 atm pressure and 310 K temperature. GaMD implemented in the GPU version of AMBER 22^12, 39^ was applied to simulate the activation of γ-secretase for cleavage of Notch. The all-atom dual-boost GaMD simulations involved an initial short cMD of 12 ns to calculate GaMD acceleration parameters and GaMD equilibration of added boost potentials for 48 ns. Three 1000 ns independent GaMD production simulations with randomized initial atomic velocities were performed on the Notch-bound γ-secretase complexes. The reference energy was set to lower bound (iE = 1) for both dihedral and total potential energy. The upper limits of the boost potential standard deviations, σ_0P_ and σ_0D_, were set to 6.0 kcal/mol for both total and dihedral potential energetic terms. The GaMD simulations are summarized in **Table S1**.

### Simulation analysis

The simulation trajectories were analyzed using VMD^40^ and CPPTRAJ^39^. The distances between Cγ atoms of catalytic aspartates PS1 residues D257 and D385, between PS1 residue D385 (atom OD2) and Notch residue V35 (atom O), between PS1 residue D385 (atom OD2) and Notch residue G34 (atom O), and between PS1 residue D385 (atom OD2) and Notch residue C33 (atom O) were calculated **(Figures S1-S4)**. The PyReweighting^13^ toolkit was applied for free energy calculations from the D257-D385 and either D385-V35, D385-G34, or D385-C33 distances for each simulation system (**Figure 1**). A bin size of 0.5 Å and cutoff of 100 frames in each bin was used to calculate the two-dimension (2D) potential mean force (PMF) free energy profiles. The hierarchical agglomerative structural clustering algorithm in CPPTRAJ^40^ was performed on GaMD simulations of cryoEM WT, cryoEM L36F, model WT, and model L36F Notch-bound γ-secretase to identify representative poses for low-energy conformational states (**Figures 2-3** and **S5-S7**). The time courses of Notch secondary structures were calculated by CPPTRAJ^40^ (**Figures S8**-**S9**).

**Figure 1.**
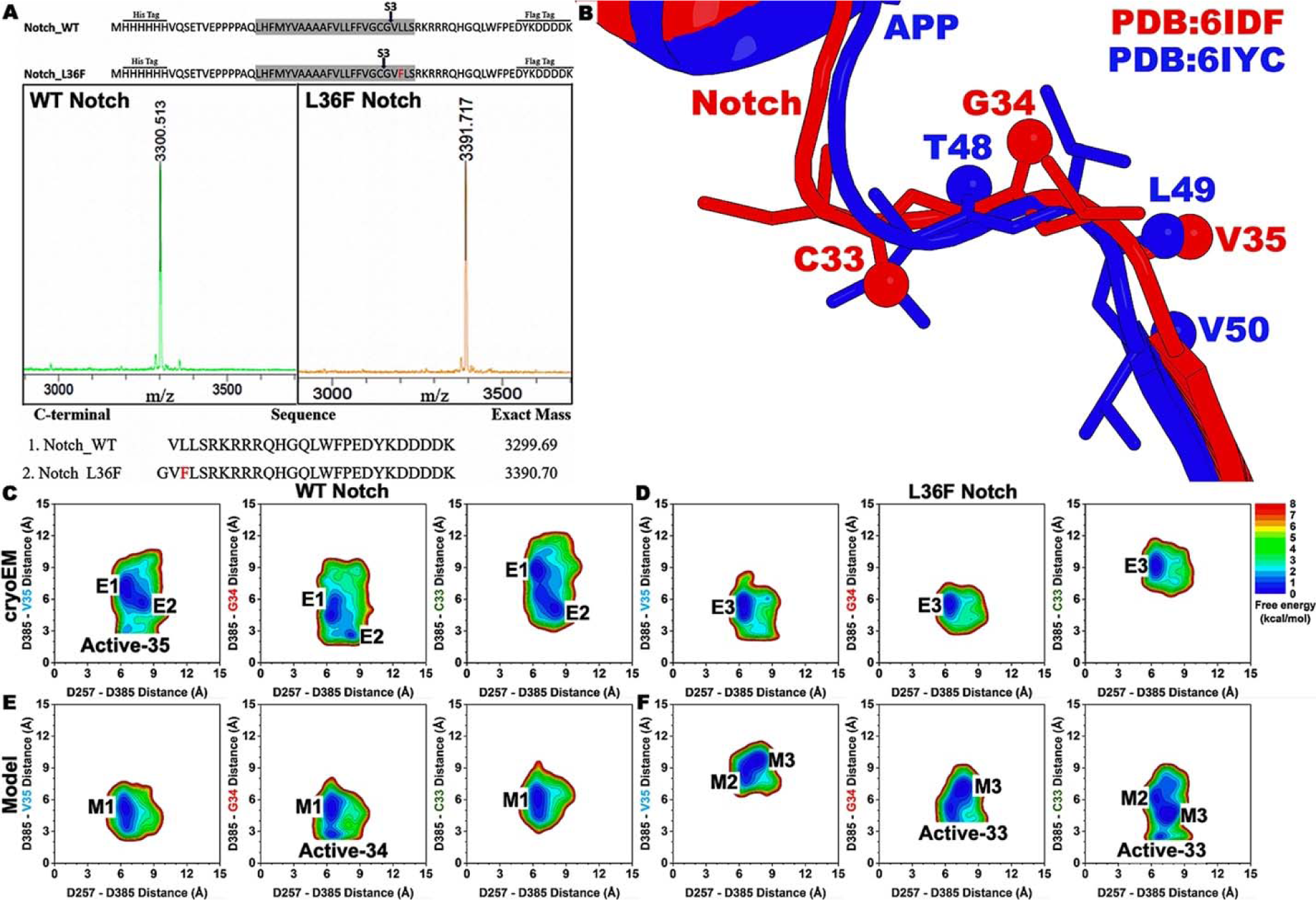
Summary of the activation of γ-secretase for cleavage of Notch. **(A)** MALDI-TOF mass spectrometric (MS) analysis of C-terminal proteolytic fragments shows shift in the S3 cleavage site between WT and L36F Notch by γ-secretase. The WT and L36F mutant sequences of Notch with observed S3 cleavage sites are shown. The transmembrane domain (TM) is highlighted in gray. N-terminal His-tag and C-terminal flag-tag are shown. MALDI-TOF MS detection of NICD-like products generated from S3 cleavage of WT and L36F variants of given Notch by γ-secretase. **(B)** Structural alignment between Notch (PDB: 6IDF) and amyloid precursor protein (APP) (PDB: 6IYC) substrates reveals an incorrect registry of the Notch substrate in the active site of γ-secretase. **(C-F)** 2D free energy profiles of the distance between PS1 residues D257 (atom Cγ) and D385 (atom Cγ) and distance between PS1 residue D385 (protonated oxygen) and either Notch residue V35, G34, or C33 (carbonyl oxygen) **(C)** in the cryoEM structure of WT Notch-bound γ-secretase (PDB: 6IDF); **(D)** in the L36F Notch-bound γ-secretase mutated from the 6IDF PDB structure; **(E)** in the model WT Notch-bound γ-secretase built from the cryoEM structure of WT APP-bound γ-secretase (PDB: 6IYC); and **(F)** in the model L36F Notch-bound γ-secretase built from the cryoEM structure of APP-bound γ-secretase (PDB: 6IYC).

**Figure 2.**
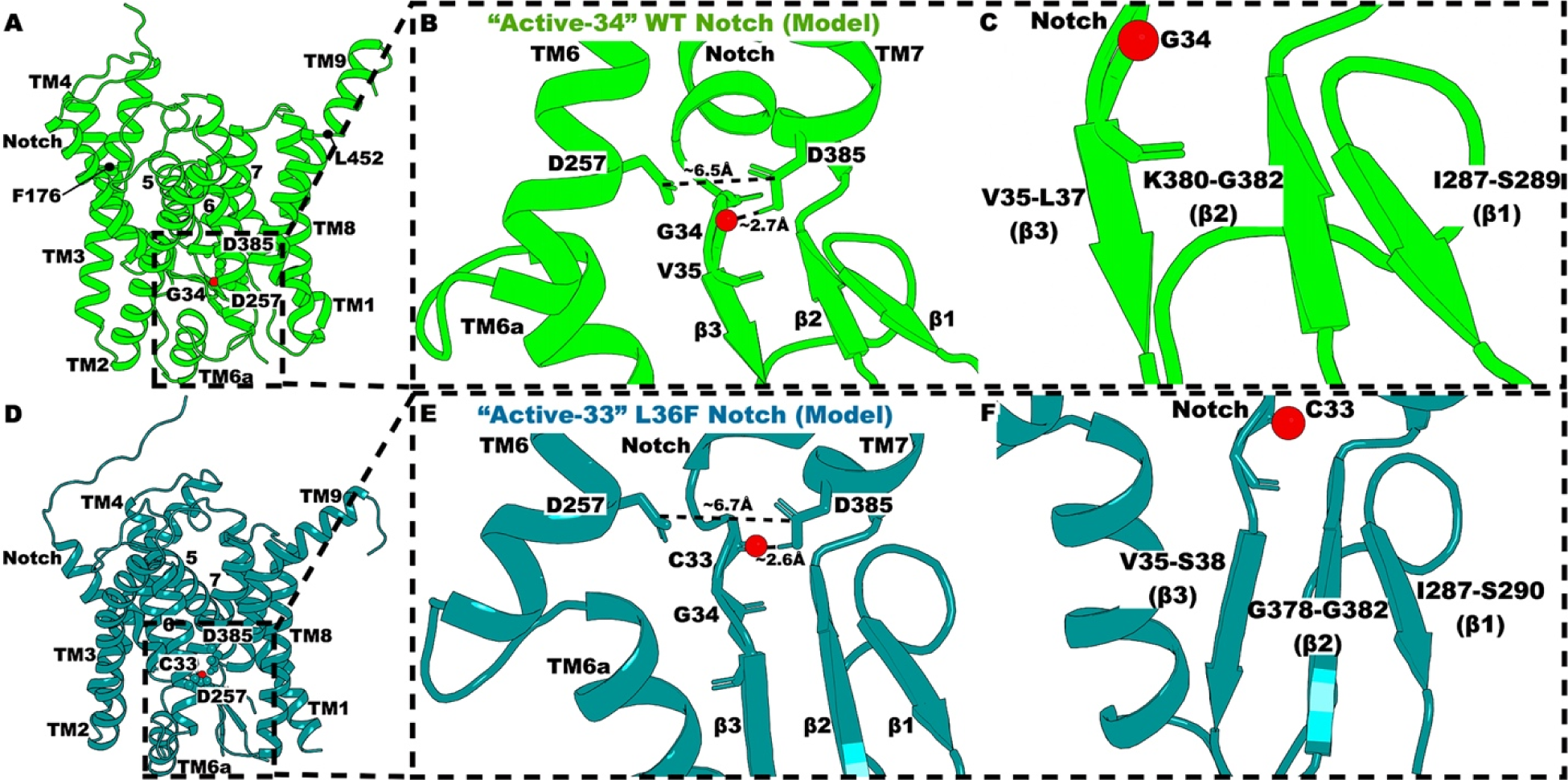
The “Active” low-energy conformational states in the model WT and L36F Notch-bound γ-secretase built from the 6IYC PDB structure. **(A)** The “Active-34” conformation of model WT Notch-bound PS1. **(B)** Active site of the model WT Notch-bound PS1 in the “Active-34” low-energy conformation. The distance between PS1 residues D257 and D385 is ∼6.5 Å, and the distance between PS1 residue D385 and Notch residue G34 is ∼2.7 Å. **(C)** Conformations of the hybrid β-sheets connected to TM6a, TM7, and Notch in the “Active-34” model WT Notch-bound PS1. **(D)** The “Active-33” conformation of model L36F Notch-bound PS1. **(E)** Active site of the model L36F Notch-bound PS1 in the “Active-33” low-energy conformation. The distance between PS1 residues D257 and D385 is ∼6.7 Å, and the distance between PS1 residue D385 and Notch residue C33 is ∼2.6 Å. **(F)** Conformations of the hybrid β-sheets connected to TM6a, TM7, and Notch in the “Active-33” model L36F Notch-bound PS1.

**Figure 3.**
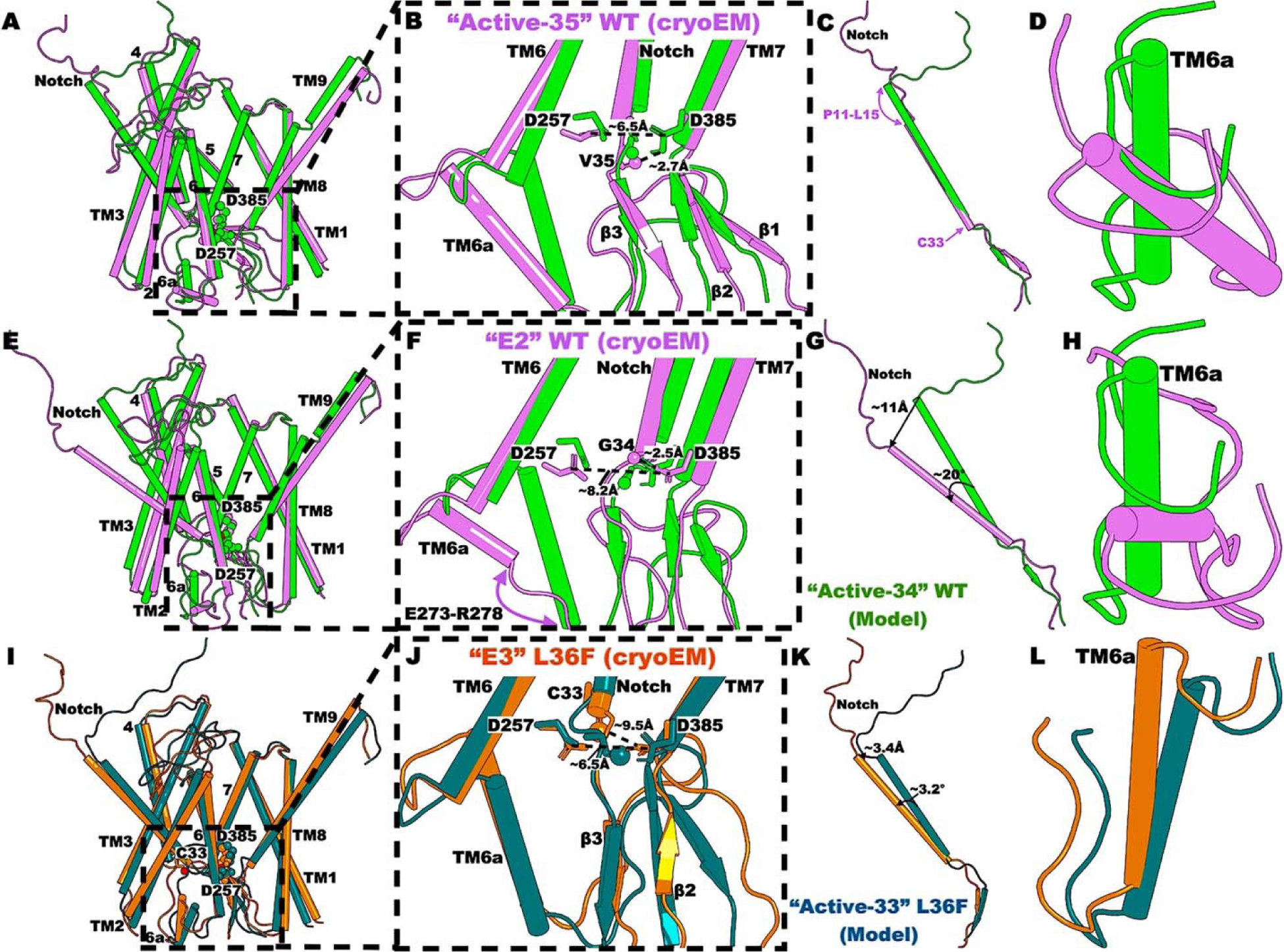
The “Active-35”, “E2”, and “E3” low-energy conformational states of cryoEM WT and L36F Notch-bound γ-secretase compared to the “Active” states of model WT and L36F Notch-bound γ-secretase. **(A)** The “Active-35” conformation of cryoEM WT Notch-bound PS1 compared to the “Active-34” conformation of model WT Notch-bound PS1. **(B)** Active site of the cryoEM WT Notch-bound PS1 in the “Active-35” low-energy conformation. The distance between PS1 residues D257 and D385 is ∼6.5 Å, and the distance between PS1 residue D385 and Notch residue V35 is ∼2.7 Å. **(C)** Conformation of the Notch substrate in the “Active-35” conformation of cryoEM WT Notch-bound PS1. **(D)** Conformation of TM6a in the “Active-35” conformation of cryoEM WT Notch-bound PS1. **(E)** The “E2” conformation of cryoEM WT Notch-bound PS1 compared to the “Active-34” conformation of model WT Notch-bound PS1. **(F)** Active site of the cryoEM WT Notch-bound PS1 in the “E2” low-energy conformation. The distance between PS1 residues D257 and D385 is ∼8.2 Å, and the distance between PS1 residue D385 and Notch residue G34 is ∼2.5 Å. **(G)** Conformation of the Notch substrate in the “E2” conformation of cryoEM WT Notch-bound PS1. **(H)** Conformation of TM6a in the “E2” conformation of cryoEM WT Notch-bound PS1. **(I)** The “E3” conformation of cryoEM L36F Notch-bound PS1 compared to the “Active-33” conformation of model L36F Notch-bound PS1. **(J)** Active site of the cryoEM L36F Notch-bound PS1 in the “E3” low-energy conformation. The distance between PS1 residues D257 and D385 is ∼6.5 Å, and the distance between PS1 residue D385 and Notch residue C33 is ∼9.5 Å. **(K)** Conformation of the Notch substrate in the “E3” conformation of cryoEM L36F Notch-bound PS1. **(L)** Conformation of TM6a in the “E3” conformation of cryoEM L36F Notch-bound PS1.

## Results and Discussion

### Cleavage of WT and L36F Notch-bound γ-secretase in biochemical experiments

We have previously shown that phenylalanine in the P2’ position in APP substrate is not tolerated by γ-secretase^25^, and leads to shifting of the cleavage site. Whether this specificity rule holds for other substrates, including Notch, is unknown. To study the effect of phenylalanine at the P2’ position on the S3 cleavage of Notch by the γ-secretase complex, enzyme reactions were performed using purified recombinant WT γ-secretase with purified recombinant Notch1-based substrates (WT or L36F; see **Figure 1A**). These substrates contained the transmembrane domain and flanking juxta-membrane residues of human Notch1, flanked on the N-terminus with a 6XHis epitope and on the C-terminus with a FLAG epitope. Enzyme reaction mixtures were then subjected to immunoprecipitation with anti-FLAG antibody for analyzing the NICD-like fragment generated after S3 cleavage by the enzyme. The immunoprecipitated product was then analyzed by matrix-assisted laser desorption ionization time-of-flight (MALDI-TOF) mass spectrometry (MS). The MALDI-TOF spectrum of NICD-like product from the enzyme reaction with WT Notch substrate shows a single cleavage product of the known G34-V35 S3 cleavage site for Notch (**Figure 1A**). In contrast, the MALDI-TOF spectrum of the NICD-like fragment from the enzyme reaction with L36F Notch shows a single product corresponding to a shifted S3 cleavage site by one amino acid in the N-terminal direction, showing cleavage at the C33-G34 site (**Figure 1A**). Thus, the intolerance of γ-secretase for phenylalanine in the P2’ position of APP substrate apparently applies to Notch substrate as well.

### Free energy profiles of cleavage of WT and L36F Notch by γ-secretase in the cryoEM systems

All-atom dual-boost GaMD simulations were carried out in parallel with biochemical experiments on the WT and L36F Notch-bound γ-secretase complexes to capture the activation of γ-secretase for cleavage of Notch (**Table S1**). GaMD simulations recorded similar averages and standard deviations of the boost potentials across the simulation systems, i.e., 14.7 ± 4.5 kcal/mol for the cryoEM WT, 14.0 ± 4.4 kcal/mol for the cryoEM L36F, 14.0 ± 4.4 kcal/mol for the model WT, and 14.9 ± 4.5 kcal/mol for the model L36F (**Table S1**). Based on our previous studies^8, 9, 20^, we chose to protonate PS1 residue D385. Furthermore, the distance between the Cγ atoms of catalytic aspartates D257 and D385 in PS1 and the distance between PS1 residue D385 (protonated oxygen) and either Notch residue V35, G34, or C33 (carbonyl oxygen) were used as reaction coordinates to calculate two-dimensional (2D) potential mean force (PMF) free energy profiles to characterize the cleavage of Notch by γ-secretase in different simulation systems (**Figure 1**). Their time courses were plotted in **Figures S1-S4**. Overall, the cleavage of Notch by γ-secretase can only be carried out when two PS1 catalytic aspartates are at a suitable distance to recruit a nucleophilic water molecule for the proteolytic reaction through water-bridged hydrogen bonds with the two aspartates^8, 9, 20^. In addition, the carbonyl group at the cleavage site on Notch (residue V35, G34, or C33) must form another hydrogen bond between its carbonyl oxygen and the protonated carboxylic side chain of catalytic aspartate D385 in PS1 for proteolysis^8, 9, 20^.

The two-dimensional (2D) potential mean force (PMF) free energy profiles of the D257-D385 distance and either D385-V35, D385-G34, or D385-C33 distance for the cryoEM WT and L36F Notch-bound γ-secretase were calculated in **Figures 1C** and **1D**, respectively. Based on these 2D free energy profiles, we failed to capture the proper cleavage of WT and L36F Notch in the cryoEM WT and L36F Notch-bound γ-secretase. In particular, the “Active” state, namely “Active-35” (cleavage at the V35-L36 site), was captured in the 2D free energy profile calculated from the D257-D385 and D385-V35 distance for the cryoEM WT Notch (**Figure 1C**). In this state, the distance between catalytic aspartates D257 and D385 in PS1 was ∼6-7 Å, while residue D385 formed a hydrogen bond with Notch residue V35 at ∼2.5-3 Å distance (**Figure 1C**). In other words, γ-secretase was activated to cleave Notch at residue V35 in the cryoEM WT Notch-bound γ-secretase, which was not the cleavage site we observed in the biochemical experiments (**Figure 1A**). Furthermore, no “Active” state was observed in the 2D free energy profile of the D257-D385 and D385-G34 distance for the cryoEM WT Notch (**Figure 1C**). We observed only the “E2” state, in which PS1 residue D385 formed a hydrogen bond with Notch residue G34 at ∼2.5-3 Å distance, but the catalytic aspartates D257 and D385 in PS1 were too far apart to recruit a nucleophilic water molecule at ∼8-8.5 Å distance (**Figure 1C**). For the cryoEM L36F simulation system, no “Active” state was observed in the 2D free energy profiles calculated from the D257-D385 distance and D385-V35, D385-G34, or D385-C33 distance (**Figure 1D**). In other words, no cleavage of L36F Notch by γ-secretase was captured in the cryoEM L36F Notch γ-secretase, which was also inconsistent with the biochemical experiments (**Figure 1A**).

Besides the “Active-35” and “E2” low-energy conformational states, we also captured the “E1” and “E3” states from the GaMD simulations of the cryoEM WT and L36F Notch-bound γ-secretase (**Figures 1C-1D**). In the “E1” low-energy conformational state, the distance between catalytic aspartates D257 and D385 was ∼6-7 Å, while the D385-V35 distance was ∼6-8 Å, D385-G34 distance was ∼4-6 Å, and D385-C33 distance was ∼8-10 Å (**Figure 1C**). In the “E3” state, while the D257-D385 and D385-C33 distance stayed at ∼6-7 Å and ∼8-10 Å, respectively, the D385-V35 and D385-G34 distance became ∼4-6.5 Å and ∼4.5-6.5 Å, respectively (**Figure 1D**).

### Structural alignment between cryoEM structures of Notch- and APP-bound γ-secretase revealed incorrect registry of Notch residues in the PS1 active site

In the effort to explain the failure to capture proper cleavage of Notch by γ-secretase in the cryoEM systems, we performed structural alignment of the substrate-bound PS1 domains in the cryoEM structures of Notch-bound γ-secretase (PDB: 6IDF)^22^ and APP-bound γ-secretase (PDB: 6IYC)^23^. Notably, we uncovered the incorrect registry of Notch residues in the PS1 active site when compared to the APP residues (**Figure 1B**). In particular, the ε-cleavage of WT APP by γ-secretase is carried out at either APP residue L49 or T48^8, 9^, cleaving the L49-V50 and T48-L49 amide bond, respectively, and the cleavage of WT Notch by γ-secretase is biochemically determined to be carried out at Notch residue G34, cleaving the G34-V35 amide bond (**Figure 1A**). However, the Notch substrate in the 6IDF^22^ PDB structure was found to be shifted one residue upwards for the proper cleavage of Notch by γ-secretase to take place (**Figure 1B**). In particular, the location of Notch residue V35 matches the location of APP residue L49 in the PS1 active site of γ-secretase (**Figure 1B**). This explained our observation that GaMD simulations captured the cleavage of WT Notch by γ-secretase at residue V35 in the cryoEM WT system (**Figure 1C**). On the other hand, the location of Notch residue G34 matched the location of APP residue T48 in the PS1 active site of γ-secretase (**Figure 1B**). This finding was consistent with a previous study by Chen and Zacharias^41^.

### Free energy profiles of cleavage of WT and L36F Notch by γ-secretase in the model systems

In order to fix the registry of Notch residues in the PS1 active site, we computationally mutated every APP residue in APP-bound γ-secretase (PDB: 6IYC)^23^ to Notch residue using the *Mutation* function of CHARMM-GUI^26-32^ as following:

APP LVFFAEDVGSNKGAIIGLMVGGVVIATVIVITLVMLKKK

Notch VQSETVEPPPPAQLHFMYVAAAAFVLLFFVGCGVLLSRK

to build the model WT Notch-bound γ-secretase. The model L36F Notch-bound γ-secretase was computationally generated from the model WT Notch system.

The 2D PMF free energy profiles of the D257-D385 distance and either D385-V35, D385-G34, or D385-C33 distance for the model WT and L36F Notch-bound γ-secretase were calculated in **Figures 1E** and **1F**, respectively. With the registry fixed, we successfully captured the correct cleavage of both WT and L36F Notch by γ-secretase in the model systems, cleaving at the G34 and C33 residues, respectively (**Figure 1A**, **1E**, and **1F**). In particular, the “Active-34” low-energy conformational state was observed in the 2D free energy profile calculated from the D257-D385 and D385-G34 distance of the model WT system (**Figure 1E**). In this state, the catalytic aspartates D257 and D385 in PS1 were close enough at ∼6-7 Å distance to recruit a nucleophilic water molecule, while PS1 residue D385 formed a proper hydrogen bond with Notch residue G34 at ∼2.5-3 Å distance (**Figure 1E**). On the other hand, the “Active-33” low-energy conformational state was observed in the 2D free energy profile calculated from the D257-D385 and D385-C33 distance of the model L36F system (**Figure 1F**). This was consistent with the experimental finding that the L36F mutation shifted cleavage site on Notch one residue upwards, from G34 (cleaving the G34-V35 amide bond) to C33 (cleaving the C33-G34 amide bond) (**Figure 1A**). In the “Active-33” state observed for the model L36F system, the distance between catalytic aspartates D257 and D385 in PS1 was ∼6-7 Å, and the D385-C33 distance was ∼2.5-3 Å (**Figure 1F**).

Besides the “Active-34” and “Active-33” low-energy conformational states captured respectively for the model WT and L36F Notch-bound γ-secretase, the “M1”, “M2”, and “M3” low-energy states were observed in the 2D free energy profiles of the model simulation systems (**Figure 1E** and **1F**). In the “M1” state, the distance between PS1 residues D257 and D385 was ∼6-7 Å, while the D385-V35, D385-G34, and D385-C33 distance were ∼4-6 Å, ∼4-6 Å, and ∼4-7 Å, respectively (**Figure 1E**). In the “M2” state, the D257-D385 distance stayed at ∼6-7 Å, while the D385-V35 and D385-C33 distance became ∼8-9.5 Å and ∼5.5-6.5 Å (**Figure 1F**). Finally, in the “M3” low-energy conformational state, the D257-D385, D385-V35, D385-G34, and D385-C33 distance were ∼7-8 Å, ∼9-10 Å, ∼6-7.5 Å, and ∼4-5 Å (**Figure 1F**).

### Active low-energy conformational states of model Notch-bound γ-secretase

The “Active-34” and “Active-33” low-energy conformational states were identified in the model WT and L36F mutant Notch-bound γ-secretase, respectively (**Figure 1E** and **1F**). They were characterized by the D257-D385 distance of ∼6-7 Å and D385-G34 or D385-C33 distance of ∼2.5-3 Å. Representative PS1 and Notch conformations of “Active-34” and “Active-33” model Notch-bound γ-secretase obtained from structural clustering of the GaMD simulation snapshots of the model WT and L36F Notch systems using CPPTRAJ^40^ were shown in **Figure 2** and compared in **Figure S5**. Overall, the Notch-bound PS1 domains of the “Active-34” WT and “Active-33” L36F were highly similar, with a C_α_-RMSD of ∼1.92 Å from structural alignment. However, it was worth noting that the helical domain of PS1 TM3 distorted at residue F176 and of PS1 TM9 distorted at residue L452 in the “Active-34” state of model WT Notch-bound PS1 (**Figure 2A**).

First, the active sites in the “Active-34” WT and “Active-33” L36F Notch-bound PS1 were shown in **Figure 2B** and **2E**, respectively. The catalytic aspartates D257 and D385 in PS1 were well positioned to recruit a nucleophilic water through water-bridged hydrogen bonds in both active states. In particular, the Cγ-atom distances between PS1 residues D257 and D385 in the “Active-34” WT and “Active-33” L36F Notch-bound PS1 were ∼6.5 Å (**Figure 2B**) and ∼6.7 Å (**Figure 2E**), respectively. Furthermore, a hydrogen bond was formed between the protonated carboxylic side chain of PS1 residue D385 and the carbonyl group of Notch residue G34 in the “Active-34” and C33 in the “Active-33”, making the carbonyl groups more electrophilic so that the proteolytic reaction can be carried out by the recruited nucleophilic water molecule. Structural alignment showed that Notch residues G34 in “Active-34” and C33 in “Active-33” state were located at similar locations in the PS1 active site (**Figure S5B**).

Second, the hybrid β-sheets connected to TM6a (β1), TM7 (β2), and Notch (β3) were different between the “Active-34” WT and “Active-33” L36F mutant Notch-bound PS1 (**Figure 2C** and **2F**). In the “Active-34” low-energy conformation, PS1 residues I287-S289 (β1) formed hybrid β-sheets with PS1 residues K380-G382 (β2) and Notch residues V35-L37 (β3) (**Figure 2C**). Meanwhile, in the “Active-33” state, the formation of antiparallel β-sheets involved PS1 residues I287-S290 (β1) and G378-G382 (β2) and Notch residues V35-S38 (β3) (**Figure 2F**).

Third, the helical domain of Notch tilted in the “Active-33” state by ∼15° compared to that in the “Active-34” state (**Figure S5C**). Notch residues P11-G32 formed the helical domain of Notch in the “Active-34” state of model WT Notch, while Notch residues Q14-G32 formed the helical domain of Notch in the “Active-33” state of model L36F mutant Notch-bound PS1. Therefore, the extracellular end of the Notch helical domain in “Active-33” state was distorted, with Notch residues P11-A13 becoming unstructured, and moved by ∼9.3 Å from that in the “Active-34” state (**Figure S5C**). Finally, the TM6a in the “Active-33” L36F mutant Notch-bound PS1 moved slightly inwards towards the PS1 active site compared to itself in the “Active-34” WT Notch-bound PS1 (**Figure S5A** and **S5D**).

### Additional low-energy conformational states of Notch-bound γ-secretase identified from GaMD simulations

In **Figure 3**, the “Active-35” and “E2” states of cryoEM WT were compared to the “Active-34” state of model WT Notch-bound PS1, while the “E3” state of cryoEM L36F mutant was compared to the “Active-33” state of model L36F mutant Notch-bound PS1. In the “Active-35” state of cryoEM WT Notch system, the two catalytic aspartates D257 and D385 in PS1 were sufficiently close at ∼6.5 Å distance to recruit a nucleophilic water molecule (**Figure 3B**). However, since Notch residue V35 in the “Active-35” state was located at the location of Notch residue G34 in the “Active-34” state of model WT Notch system, the protonated D385 formed a hydrogen bond with Notch residue V35 at ∼2.7 Å distance, which led to the improper cleavage of Notch by γ-secretase observed for the cryoEM WT Notch system (**Figures 3B** and **1C**). Antiparallel hybrid β-sheets were formed between PS1 residues I287-S290 (β1) and V379-G382 (β2) and Notch residues L36-R39 (β3) in the “Active-35” low-energy state of the cryoEM WT Notch-bound γ-secretase (**Figure 3B**). Furthermore, the helical domain of Notch also varied between the “Active-35” state of cryoEM WT Notch and “Active-34” state of model WT Notch system (**Figure 3C**). In particular, the extracellular end of Notch helical domain in the “Active-35” state was distorted at Notch residues P11-L15, while Notch residue C33 in the “Active-35” state turned helical compared to the “Active-34” low-energy conformation (**Figure 3C**). Lastly, the TM6a in the “Active-35” state of cryoEM WT Notch tilted significantly compared to that in the “Active-34” state of model WT Notch-bound γ-secretase (**Figure 3D**).

In the “E2” low-energy conformational state of cryoEM WT Notch-bound γ-secretase, the protonated D385 formed a hydrogen bond with the carbonyl oxygen of Notch residue G34 at ∼2.5 Å distance (**Figure 3F**). However, due to the incorrect registry of Notch residue G34 in the PS1 active site, the catalytic aspartates D257 and D385 in PS1 were too far apart at ∼8.2 Å distance to form water-bridged hydrogen bonds to recruit a nucleophilic water molecule (**Figure 3F**). The hybrid β-sheets were all distorted in the “E2” state of cryoEM WT Notch-bound γ-secretase (**Figure 3F**). Furthermore, the extracellular end of Notch helical domain in the “E2” state tilted by ∼20° and moved by ∼11 Å compared to that in the “Active-34” state of model WT Notch-bound PS1 (**Figure 3G**). Finally, the TM6a in the “E2” state titled significantly compared to the “Active-34” state, and the TM6a helical domain was distorted at PS1 residues E273-R278 in the “E2” state of cryoEM WT Notch-bound γ-secretase (**Figure 3F** and **3H**).

Overall, the “E1” state can be referred to as the intermediate between the “Active-35” and “E2” state of the cryoEM WT Notch-bound γ-secretase (**Figure S6A-D**). In particular, similar to the “E2” state, the hybrid β-sheets were completely distorted in the “E1” low-energy conformational state (**Figure S6B**). However, like the “Active-35” low-energy conformational state, the extracellular end of Notch helical domain was distorted at Notch residues P11-A13, while Notch residue C33 turned helical in the “E1” state of cryoEM WT Notch-bound γ-secretase (**Figure S6C**).

In the “E3” low-energy conformational state of cryoEM L36F mutant Notch system, the Cγ-atom distance between PS1 residues D257 and D385 was ∼6.5 Å (**Figure 3J**). However, the incorrect registry of the Notch substrate in the PS1 active site made Notch residue C33 too far apart from the protonated D385 at ∼9.5 Å distance (**Figure 3J**). The hybrid β-sheets were only formed between PS1 residues V379-L381 (β2) and Notch residues L37-R39 (β3), while the β1 domain became unstructured in the “E3” low-energy state (**Figure 3J**). Furthermore, the extracellular end of Notch helical domain in the “E3” state of cryoEM L36F mutant Notch-bound PS1 titled by ∼3.2 Å and only moved by ∼3.4 Å compared to that in the “Active-33” model L36F mutant Notch-bound γ-secretase (**Figure 3K**).

The “M1” low-energy conformational state of model WT Notch-bound γ-secretase was compared to the “Active-34” state in **Figure S6E-S6H**, while the “M2” and “M3” low-energy conformations of model L36F mutant Notch-bound γ-secretase was compared to the “Active-33” state in **Figure S7**. Overall, the PS1 active site in the “M1” state was highly similar to that in the “Active-34” state of model WT Notch-bound PS1 in terms of the D257-D385 distance and residues involved in the hybrid sheet formation (**Figure S6F**). However, in the “M1” state, the carbonyl group of Notch residue G34 flipped towards PS1 residue D257, resulting in the D385-G34 distance of ∼5.0 Å (**Figure S6F**). The flipping towards the other catalytic aspartate of the carbonyl group of Notch residue G34 opened the possibility of forming a hydrogen bond with PS1 residue D257 and thus the proton exchange between PS1 residues D257 and D385. This was consistent with previous studies where we observed the activation of γ-secretase for cleavage of APP with either protonation (protonated D257^8^ or D385^9^). Notably, PS1 residues F105, Y106, and T107 of the hydrophilic loop 1 (HL1) turned α-helical in the “M2” and “M3” state of model L36F mutant Notch-bound γ-secretase (**Figure S7A** and **S7E**), while remained unstructured in the other low-energy conformations. Residue Y106 of HL1 was experimentally shown to play an important role in the binding of γ-secretase modulators (GSMs)^42^. Furthermore, the β3-domain was completely distorted in the “M2” low-energy state compared to the “Active-33” and “M3” states of model L36F mutant Notch-bound γ-secretase (**Figure S7B**). Finally, the extracellular end of Notch helical domain was distorted at Notch residues P11-L15 in both the “M2” and “M3” states compared to the “Active-33” low-energy conformation of model L36F mutant Notch-bound γ-secretase (**Figure S7C** and **S7G**).

### Secondary structures of Notch substrate in the WT and L36F mutant Notch-bound γ-secretase

Time courses of Notch secondary structures for the model WT and L36F mutant Notch-bound γ-secretase are shown in **Figure S8**, while time courses of Notch secondary structures for the cryoEM WT and L36F mutant Notch-bound γ-secretase are shown in **Figure S9**. Overall, we observed β-sheet secondary structure in the C-terminus of Notch (β3) in all the simulations of both model and cryoEM Notch-bound γ-secretase. In the model WT and L36F mutant systems, the formation of the β3-domain mostly involved Notch residues L36-S38 (**Figure S8**). Notably, even though a shift in cleavage site from Notch residue G34 to C33 was observed from the WT to L36F mutant system, the residues forming β3-domain did not change between the two systems (**Figures S8** and **2**). In the cryoEM WT simulation system, the Notch substrate was improperly cleaved at residue V35 by γ-secretase (**Figures 1C** and **3B**). In the cryoEM L36F mutant simulation system, no cleavage of the Notch substrate γ-secretase was observed in the GaMD simulations (**Figure 1D** and **3J**). Nevertheless, the β3 β-sheet was still formed at Notch residues L37-R39 in the cryoEM WT and L36F mutant Notch-bound γ-secretase for most of the simulations (**Figure S9**). Only in Sim3 of the cryoEM L36F simulation system, the β3 β-sheet was shortened, with Notch residue R39 being turned or unstructured for most of the simulation (**Figure S9F**). These findings have shown that the formation of the β3 β-sheet may not be the primary factor for γ-secretase to determine the cleavage site on Notch or other substrates^8, 9, 20^. This was consistent with our previous study where we observed the cleavage of the Aβ49 peptide by γ-secretase even when the C-terminus of the substrate remained unstructured^20^.

In the model WT Notch simulation system, the α-helical domain of Notch mostly consisted of Notch residues P11-G32 (**Figure S8A**-**S8C**). However, Notch residue L27 usually existed as turn or 3-10-helix, while residues G32-C33 occasionally became turn during the simulations (**Figure S8A-S8C**). The length of the α-helical domain of Notch significantly reduced in the model L36F mutant simulation system compared to the model WT Notch (**Figure S8D-S8F**). In particular, the α-helical domain of Notch in the model L36F mutant system only spanned from Notch residue H16 to V26 (**Figure S8D-S8F**). Notch residue L27 mostly existed as turn in the L36F mutation, except for the last 800 ns of Sim3 (**Figure S8F**) where it exhibited α-helical secondary structure. Notch residues L28-V31 mostly showed 3-10-helical secondary structure, except for the last 200ns of Sim3 of the L36F mutation system, where they turned α-helical (**Figure S8F**). Notch residue G32 (right next to the cleavage site of the L36F mutant Notch) primarily existed as turn in all three simulations (**Figure S8D-S8F**).

The α-helical domain of Notch did not exhibit significant changes from the cryoEM WT to L36F mutant Notch-bound γ-secretase (**Figure S9**). In particular, the α-helical domain of Notch primarily spanned from Notch residue Q14 to G32 in both simulation systems (**Figure S9**). The decrease in the length of Notch α-helical domain as well as the tilted and distorted TM6a (**Figures S9, S6, S7,** and **3**) may have shown a decrease in structural stability from the model to cryoEM Notch-bound γ-secretase. Furthermore, Notch residues C33-G34 mostly existed as turn in the GaMD simulations of both cryoEM WT and L36F mutant Notch-bound γ-secretase (**Figure S9**). In the cryoEM WT simulation system, Notch residues V35-L36 also existed as turn, especially in Sim2 and Sim3 (**Figure S9B-S9C**).

## Conclusions

We have successfully captured the dynamic activation of γ-secretase for cleavage of WT and L36F mutant Notch for the first time by performing GaMD simulations alongside mass spectrometry (MS) experiments. Through the MALDI-TOF spectra, γ-secretase was experimentally determined to cleave WT Notch at residue G34 (cleaving the G34-V35 amide bond, consistent with an earlier report^43^ and L36F Notch at residue C33 (cleaving the C33-G34 amide bond) (**Figure 1A**). Initially, we used the 6IDF PDB^22^ structure of Notch-bound γ-secretase to simulate γ-secretase activation for cleavage of Notch. However, we failed to capture the proper cleavage for WT Notch at residue G34 and cleavage of L36F Notch by γ-secretase (**Figure 1C-1D**). We then uncovered an incorrect registry of the Notch substrate in the PS1 active site by structurally aligning the PDB structures of Notch-bound (PDB: 6IDF^22^) and APP-bound γ-secretase (PDB: 6IYC^23^), with which we have successfully capture γ-secretase activation for proper cleavage of APP with either protonation of catalytic aspartates (PS1 residue D257^8^ or D385^9^). The finding was consistent with a previous study by Chen and Zacharias^41^. We then mutated every residue of APP to Notch residue to prepare model Notch-bound γ-secretase complexes and successfully captured the correct cleavages of WT and L36F Notch at Notch residue G34 and C33, respectively, by γ-secretase in our GaMD simulations (**Figure 1E-1F**). The shift of cleavage site in the L36F-like mutation of Notch by γ-secretase has been observed in our previous study of the analogous M51F mutant APP-bound γ-secretase^8^.

Similar to the mechanism of γ-secretase activation for cleavage of APP and Aβ peptides uncovered from our previous studies^8, 9, 20^, two conditions must be met for proper cleavage of Notch by γ-secretase (**Figure 2**). First, the two catalytic aspartates D257 and D385 in PS1 must be sufficiently close, at ∼6-7 Å distance, to recruit a nucleophilic water molecule through water-bridged hydrogen bonds. Second, the carbonyl group of Notch residue at Notch cleavage site must form a hydrogen bond with the protonated carboxylic side chain of PS1 residue D385 to make the carbonyl group more electrophilic for the proteolytic reaction to take place (**Figure 2**). In most previous studies by other groups, only one of the listed conditions here was shown in the simulation analysis results, which resulted from using apo γ-secretase structure without any substrate^44^, focusing on only the catalytic aspartate-substrate residue interaction^41^, etc.

Furthermore, by analyzing the low-energy conformational states and Notch secondary structures during the GaMD simulations, we proposed that the β3 β-sheet might not be the critical factor for γ-secretase to determine the proper cleavage site on Notch or other substrates^8, 9, 20^. This was consistent with our previous study where we observed the cleavage of the Aβ49 peptide by γ-secretase even when the C-terminus of the substrate remained unstructured^20^. Moreover, due to the distorted α-helical domain in the Notch substrate and TM6a observed in our GaMD simulation, we suggest that the cryoEM structure of Notch-bound γ-secretase (PDB: 6IDF^22^) deviates from activation-compatible conformation of the enzyme-substrate complex as the model Notch-bound γ-secretase built from the cryoEM structure of APP-bound γ-secretase (PDB: 6IYC^23^). Finally, since the carbonyl group of Notch residue G34 could flip towards either catalytic aspartate (**Figures 2B** and **S6F**), we propose that proton exchange very likely takes place between the two catalytic aspartates. This is consistent with our ability to capture γ-secretase activation for cleavage of APP with either catalytic aspartate protonated in previous studies^8, 9^. Our findings here provided mechanistic insights into the structural dynamics and enzyme-substrate interactions required for γ-secretase activation for cleavage of Notch and other substrates.

## Acknowledgements

This work used supercomputing resources with allocation award TG-MCB180049 through the Extreme Science and Engineering Discovery Environment (XSEDE), which is supported by National Science Foundation grant number ACI-1548562, and project M2874 through the National Energy Research Scientific Computing Center (NERSC), which is a U.S. Department of Energy Office of Science User Facility operated under Contract No. DE-AC02-05CH11231, and the Research Computing Cluster at the University of Kansas. This work was supported in part by the startup funding in the College of Liberal Arts and Sciences at the University of Kansas and award 2121063 from National Science Foundation (Y.M.) and AG66986 from the National Institutes of Health (M.S.W.).

## Author Contributions

H.N.D. performed GaMD simulations, analyzed simulation data and wrote the manuscript. S.R.M. performed enzyme reactions, analyzed mass spectrometry results and wrote the manuscript. Y.M. and M.S.W. supervised the project, interpreted data, and wrote the manuscript. All authors contributed towards the final version of the manuscript.

## Competing Interests

The authors declare no competing interests.

## Notes

### Competing Interest Statement

The authors have declared no competing interest.

## References

1. Kopan, R.; Ilagan, M. X. G. γ-Secretase: proteasome of the membrane? Nature Reviews Molecular Cell Biology 2004, 5, 499–504.

2. Wolfe, M. S.; Miao, Y. Structure and mechanism of the gamma-secretase intramembrane protease complex. Current Opinion in Structural Biology 2022, 74, 102373.

3. Bai, X. C.; Yan, C. Y.; Yang, G. H.; Lu, P. L.; Ma, D.; Sun, L. F.; Zhou, R.; Scheres, S. H. W.; Shi, Y. G. An atomic structure of human gamma-secretase. Nature 2015, 525 (7568), 212–217. DOI: 10.1038/nature14892.

4. Wolfe, M. S.; Xia, W.; Ostaszewski, B. L.; Diehl, T. S.; Kimberly, W. T.; Selkoe, D. J. Two transmembrane aspartates in presenilin-1 required for presenilin endoproteolysis and gamma-secretase activity. Nature 1999, 398, 513–517.

5. Wolfe, M. Structure and Function of the gamma-Secretase Complex. Biochemistry 2019, 58 (27), 2953.

6. Guner, G.; Lichtenthaler, S. F. The substrate repertoire of gamma-secretase/presenilin. Seminars in Cell & Developmental Biology 2020, 105, 27–42.

7. Dehury, B.; Somavarapu, A.; Kepp, K. A computer-simulated mechanism of familial Alzheimer’s disease: Mutations enhance thermal dynamics and favor looser substrate-binding to gamma-secretase. Journal of Structural Biology 2020, 212, 107648.

8. Bhattarai, A.; Devkota, S.; Bhattarai, S.; Wolfe, M. S.; Miao, Y. Mechanisms of gamma-secretase activation and substrate processing. ACS Central Science 2020, 6 (6), 969–983.

9. Do, H. N.; Devkota, S.; Bhattarai, A.; Wolfe, M. S.; Miao, Y. Effects of presenilin-1 familial Alzheimer’s disease mutations on γ-secretase activation for cleavage of amyloid precursor protein. Communications Biology 2023, 6 (174).

10. Karplus, M.; McCammon, J. A. Molecular dynamics simulations of biomolecules. Nature Structural and Molecular Biology 2002, 9, 646–652.

11. Wang, J.; Arantes, P.; Bhattarai, A.; Hsu, R.; Pawnikar, S.; Huang, Y.-m.; Palermo, G.; Miao, Y. Gaussian accelerated molecular dynamics: principles and applications. WIREs Computational Molecular Science 2021, e1521. DOI: 10.1002/wcms.1521.

12. Miao, Y.; Feher, V. A.; McCammon, J. A. Gaussian accelerated molecular dynamics: unconstrained enhanced sampling and free energy calculation. Journal of Chemical Theory and Computation 2015, 11, 3584–3595.

13. Miao, Y.; Sinko, W.; Pierce, L.; Bucher, D.; Walker, R. C.; McCammon, J. A. Improved reweighting of accelerated molecular dynamics simulations for free energy calculation. Journal of Chemical Theory and Computation 2014, 10, 2677–2689.

14. Miao, Y.; Bhattarai, A.; Wang, J. Ligand Gaussian accelerated molecular dynamics (LiGaMD): characterization of ligand binding thermodynamics and kinetics. Journal of Chemical Theory and Computation 2020, 16, 5526–5547.

15. Wang, J.; Miao, Y. Peptide Gaussian accelerated molecular dynamics (Pep-GaMD): enhanced sampling and free energy and kinetics calculations of peptide binding. Journal of Chemical Physics 2020, 153, 154109.

16. Wang, J.; Miao, Y. Protein-Protein Interaction-Gaussian Accelerated Molecular Dynamics (PPI-GaMD): Characterization of Protein Binding Thermodynamics and Kinetics. Journal of Chemical Theory and Computation 2022, 18 (3), 1275–1285.

17. Wang, J.; Miao, Y. Ligand Gaussian Accelerated Molecular Dynamics 2 (LiGaMD2): Improved Calculations of Ligand Binding Thermodynamics and Kinetics with Closed Protein Pocket. Journal of Chemical Theory and Computation 2023, 19 (3), 733–745.

18. Do, H. N.; Wang, J.; Bhattarai, A.; Miao, Y. GLOW: A Workflow Integrating Gaussian-Accelerated Molecular Dynamics and Deep Learning for Free Energy Profiling. Journal of Chemical Theory and Computation 2022, 18 (3), 1423–1436.

19. Do, H. N.; Miao, Y. Deep Boosted Molecular Dynamics (DBMD): Accelerating Molecular Simulations with Gaussian Boost Potentials Generated Using Probabilistic Bayesian Deep Neural Network. The Journal of Physical Chemistry Letters 2023, 14 (21), 4970–4982. DOI: 10.1021/acs.jpclett.3c00926.

20. Bhattarai, A.; Devkota, S.; Do, H.; Wang, J.; Bhattarai, S.; Wolfe, M.; Miao, Y. Mechanism of Tripeptide Trimming of Amyloid beta-Peptide 49 by gamma-Secretase. Journal of American Chemical Society 2022, 144, 6215–6226.

21. Takami, M.; Nagashima, Y.; Sano, Y.; Ishihara, S.; Morishima-Kawashima, M.; Funamoto, S.; Ihara, Y. γ-Secretase: Successive Tripeptide and Tetrapeptide Release from the Transmembrane Domain of β-Carboxyl Terminal Fragment. Journal of Neuroscience 2009, 29 (41), 13042–13052.

22. Yang, G.; Zhou, R.; Zhou, Q.; Guo, X.; Yan, C.; Ke, M.; Lei, J.; Shi, Y. Structural basis of Notch recognition by human gamma-secretase. Nature 2019, 565, 192–197.

23. Zhou, R.; Yang, G.; Guo, X.; Zhou, Q.; Lei, J.; Shi, Y. Recognition of the amyloid precursor protein by human gamma-secretase. Science 2019, 363 (6428), aaw0930.

24. Lu, P.; Bai, X.-C.; Ma, D.; Xie, T.; Yan, C.; Sun, L.; Yang, G.; Zhao, Y.; Zhou, R.; Scheres, S. H.;, et al. Three-dimensional structure of human gamma-secretase. Nature 2014, 512, 166–170.

25. Bolduc, D. M.; Montagna, D. R.; Seghers, M. C.; Wolfe, M. S.; Selkoe, D. J. The amyloid-beta forming tripeptide cleavage mechanism of gamma-secretase. eLife 2016, 17578.

26. Jo, S.; Kim, T.; Iyer, V.; Im, W. CHARMM-GUI: A Web-based Graphical User Interface for CHARMM. Journal of Computational Chemistry 2008, 29, 1859–1865.

27. Brooks, B.; Brooks III, C.; MacKerell Jr, A.; Nilsson, L.; Petrella, R.; Roux, B.; Won, Y.; Archontis, G.; Bartels, C.; Boresch, S.;, et al. CHARMM: The Biomolecular Simulation Program. Journal of Computational Chemistry 2009, 30, 1545–1614.

28. Lee, J.; Cheng, X.; Swails, J.; Yeom, M.; Eastman, P.; Lemkul, J.; Wei, S.; Buckner, J.; Jeong, J.; Qi, Y.;, et al. CHARMM-GUI Input Generator for NAMD, GROMACS, AMBER, OpenMM, and CHARMM/OpenMM Simulations using the CHARMM36 Additive Force Field. Journal of Chemical Theory and Computation 2016, 12, 405-413.

29. Wu, E.; Cheng, X.; Jo, S.; Rui, H.; Song, K.; Davila-Contreras, E.; Qi, Y.; Lee, J.; Monje-Galvan, V.; Venable, R.;, et al. CHARMM-GUI Membrane Builder Toward Realistic Biological Membrane Simulations. Journal of Computational Chemistry 2014, 35, 1997–2004.

30. Jo, S.; Kim, T.; Im, W. Automated Builder and Database of Protein/Membrane Complexes for Molecular Dynamics Simulations. PLoS ONE 2007, 2 (9), e880.

31. Lee, J.; Patel, D.; Stahle, J.; Park, S.-J.; Kern, N.; Kim, S.; Lee, J.; Cheng, X.; Valvano, M.; Holst, O.;, et al. CHARMM-GUI Membrane Builder for Complex Biological Membrane Simulations with Glycolipids and Lipoglycans. Journal of Chemical Theory and Computation 2019, 15, 775–786.

32. Lee, J.; Hitzenberger, M.; M. Rieger; N.R. Kern; M. Zacharias; Im, W. CHARMM-GUI supports the Amber force fields. Journal of Chemical Physics 2020, 153, 035103.

33. Huang, J.; Rauscher, S.; Nawrocki, G.; Ran, T.; Feig, M.; de Groot, B. L.; Grubmuller, H.; MacKerell Jr, A. D. CHARMM36m: an improved force field for folded and intrinscially disordered proteins. Nature Methods 2017, 14, 71–73.

34. Ryckaert, J.-P.; Ciccotti, G.; Berendsen, H. J. C. Numerical integration of the cartesian equations of motion of a system with constraints: molecular dynamics of n-alkanes. Journal of Computational Physics 1977, 23 (3), 327–341.

35. Evans, D. J. Computer “experiment” for nonlinear thermodynamics of Couette flow. The Journal of Chemical Physics 1983, 78, 3297.

36. Hoover, W. G.; Ladd, A. J. C.; Moran, B. High-Strain-Rate Plastic Flow Studied via Nonequilibrium Molecular Dynamics. Physical Review Letters 1982, 48, 1818.

37. Berendsen, H. J. C.; Postma, J. P. M.; Vangunsteren, W. F.; Dinola, A.; Haak, J. R. Molecular Dynamics with Coupling to an External Bath. Journal of Chemical Physics 1984, 81 (8), 3684–3690.

38. Essmann, U.; Perera, L.; Berkowitz, M. L.; Darden, T.; Lee, H.; Pedersen, L. G. A Smooth Particle Mesh Ewald Method. Journal of Chemical Physics 1995, 103 (19).

39. Case, D. A.; Aktulga, H. M.; Belfon, K.; Ben-Shalom, I. Y.; Berryman, J. T.; Brozell, S. R.; Cerutti, D. S.; Cheatham, I., T.E.; Cisneros, G. A.; Cruzeiro, V. W. D.;, et al. Amber 2023. University of California, San Francisco 2023.

40. Roe, D. R.; Cheatham, I. T. E. PTRAJ and CPPTRAJ: software for processing and analysis of molecular dynamics trajectory data. Journal of Chemical Theory and Computation 2013, 9, 3084–3095.

41. Chen, S.-Y.; Zacharias, M. An internal docking site stabilizes substrate binding to γ-secretase: Analysis by molecular dynamics simulations Biophysical Journal 2022, 121 (12), 2330–2344.

42. Liu, L.; Lauro, B. M.; Wolfe, M. S.; Selkoe, D. J. Hydrophilic loop 1 of Presenilin-1 and the APP GxxxG transmembrane motif regulate gamma-secretase function in generating Alzheimer-causing Abeta peptides. Journal of Biological Chemistry 2021, 296, 100393.

43. Schroeter, E. H.; Kisslinger, J. A.; Kopan, R. Notch-1 signaling requires ligand-induced proteolytic release of intracellular domain. Nature 1998, 393 (6683), 382–386.

44. Aguayo-Ortiz, R.; Chávez-García, C.; Straub, J. E.; Dominguez, L. Characterizing the structural ensemble of γ-secretase using a multiscale molecular dynamics approach. Chemical Science 2017, 8, 5576–5584.

