## Supporting Information for "Molecular Dynamics Activation of γ-Secretase for Cleavage of Notch1 Substrate"

**Figure S1**. Time courses of distances between PS1 residues D257 (atom Cγ) and D385 (atom atom Cγ) **(A)**, PS1 residue D385 (protonated oxygen) and Notch residue V35 (carbonyl oxygen) **(B)**, PS1 residue D385 (protonated oxygen) and Notch residue G34 (carbonyl oxygen) **(C)**, and PS1 residue D385 (protonated oxygen) and Notch residue C33 (carbonyl oxygen) **(D)** in the cryoEM WT Notch-bound γ-secretase (PDB: 6IDF).

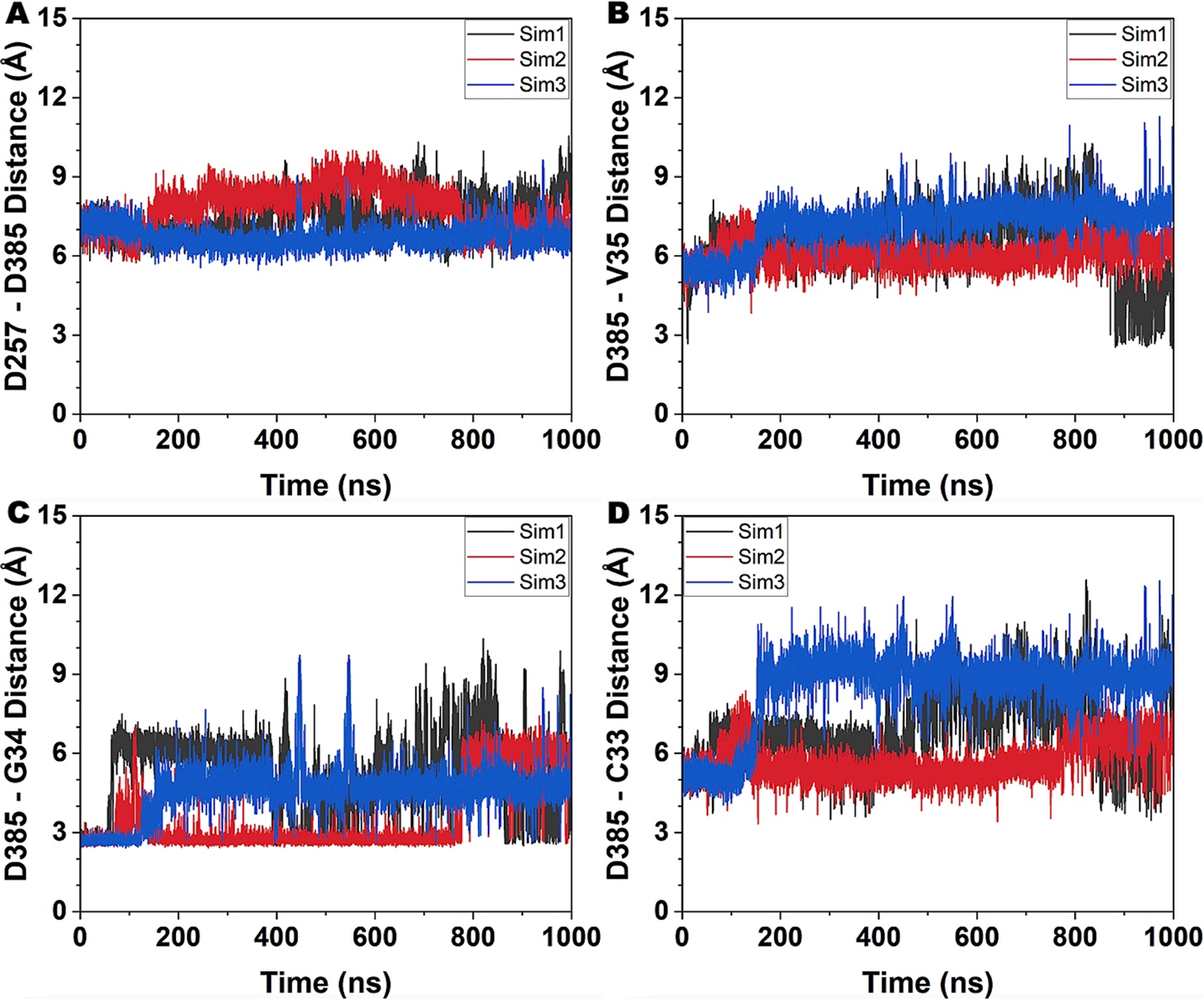

**Figure S2**. Time courses of distances between PS1 residues D257 (atom Cγ) and D385 (atom atom Cγ) **(A)**, PS1 residue D385 (protonated oxygen) and Notch residue V35 (carbonyl oxygen) **(B)**, PS1 residue D385 (protonated oxygen) and Notch residue G34 (carbonyl oxygen) **(C)**, and PS1 residue D385 (protonated oxygen) and Notch residue C33 (carbonyl oxygen) **(D)** in the cryoEM L36F Notch-bound γ-secretase mutated from the 6IDF PDB structure.

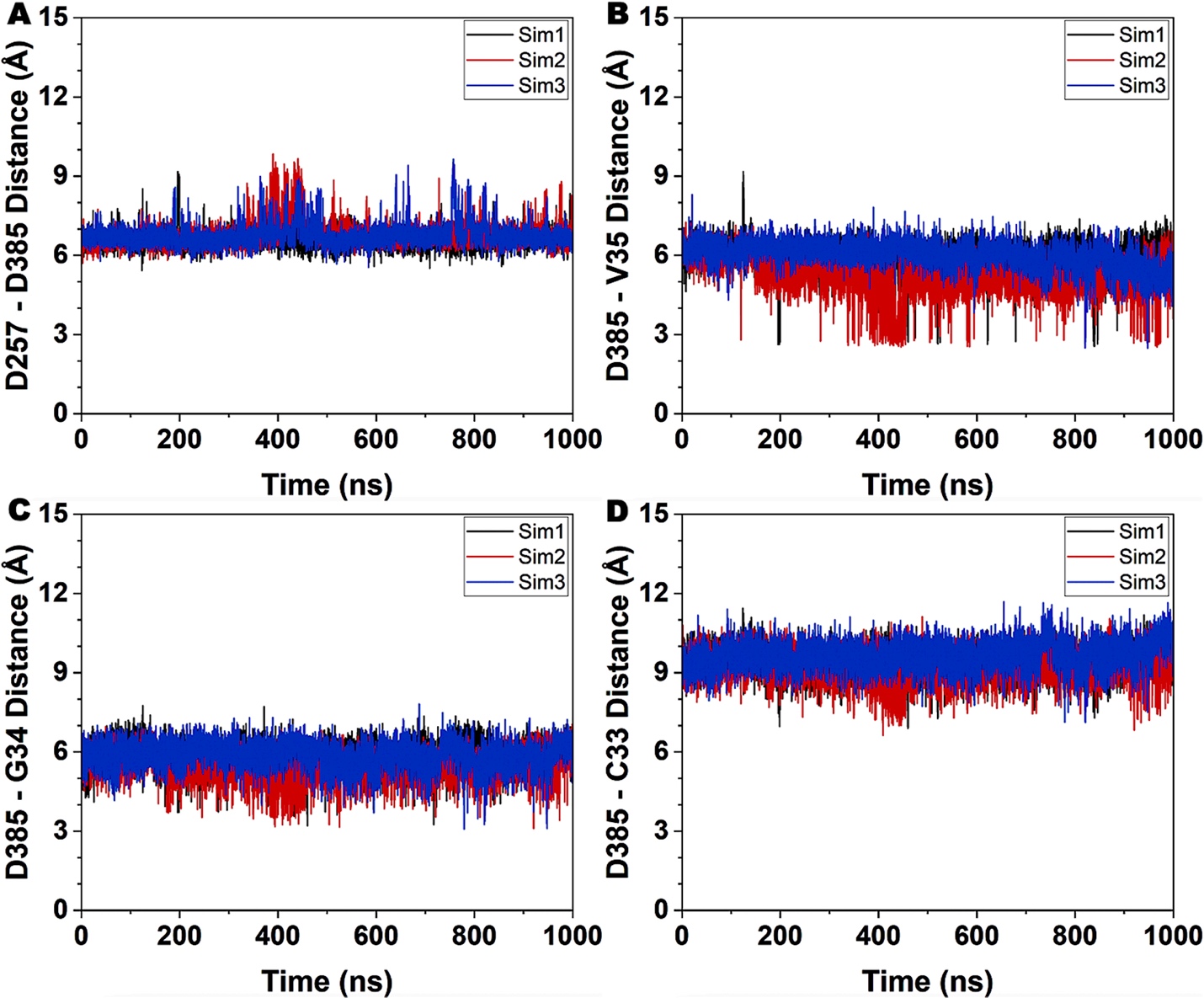

**Figure S3**. Time courses of distances between PS1 residues D257 (atom Cγ) and D385 (atom atom Cγ) **(A)**, PS1 residue D385 (protonated oxygen) and Notch residue V35 (carbonyl oxygen) **(B)**, PS1 residue D385 (protonated oxygen) and Notch residue G34 (carbonyl oxygen) **(C)**, and PS1 residue D385 (protonated oxygen) and Notch residue C33 (carbonyl oxygen) **(D)** in the model WT Notch-bound γ-secretase built from the 6IYC PDB structure.

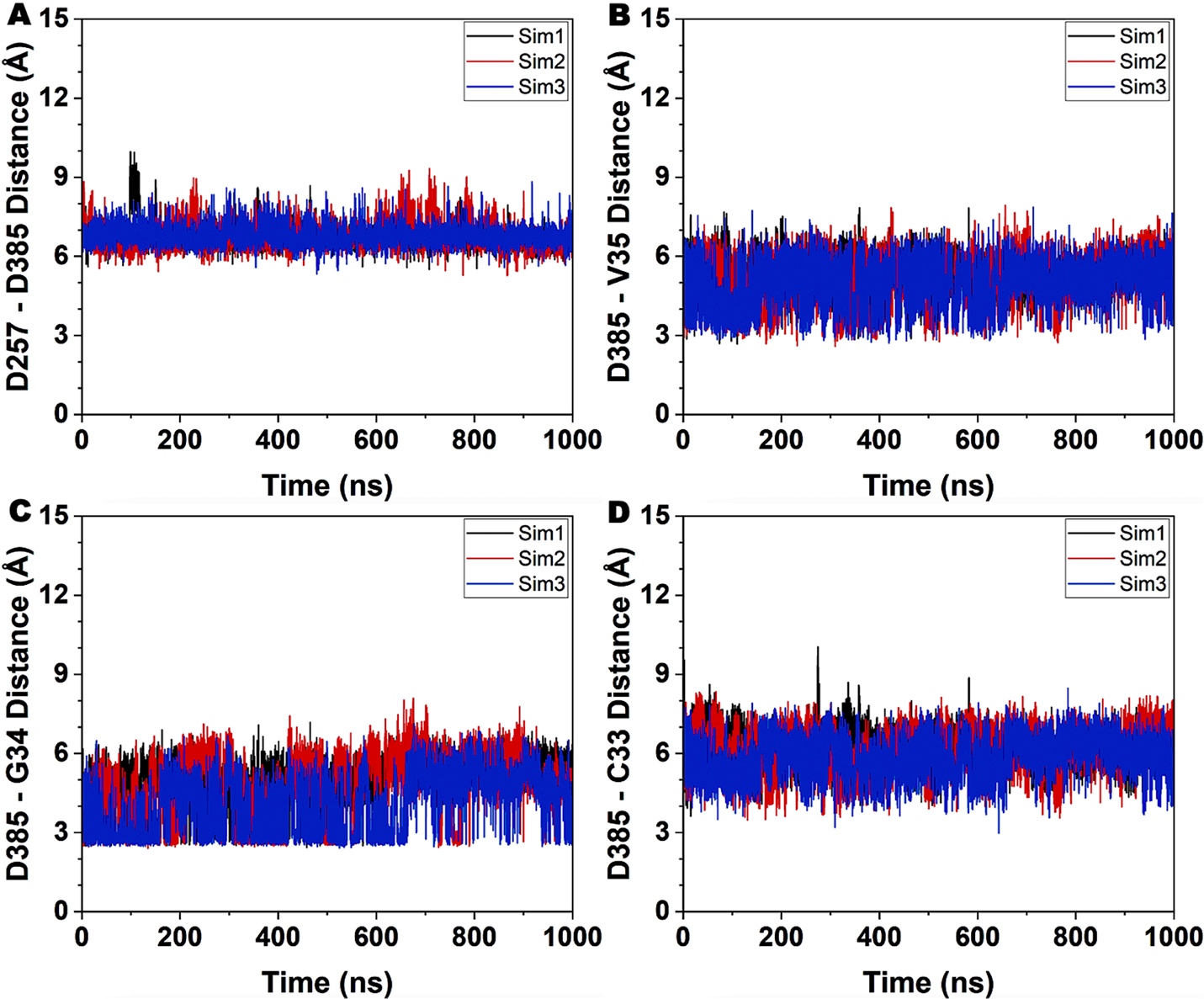

**Figure S4**. Time courses of distances between PS1 residues D257 (atom Cγ) and D385 (atom atom Cγ) **(A)**, PS1 residue D385 (protonated oxygen) and Notch residue V35 (carbonyl oxygen) **(B)**, PS1 residue D385 (protonated oxygen) and Notch residue G34 (carbonyl oxygen) **(C)**, and PS1 residue D385 (protonated oxygen) and Notch residue C33 (carbonyl oxygen) **(D)** in the model L36F Notch-bound γ-secretase built from the 6IYC PDB structure.

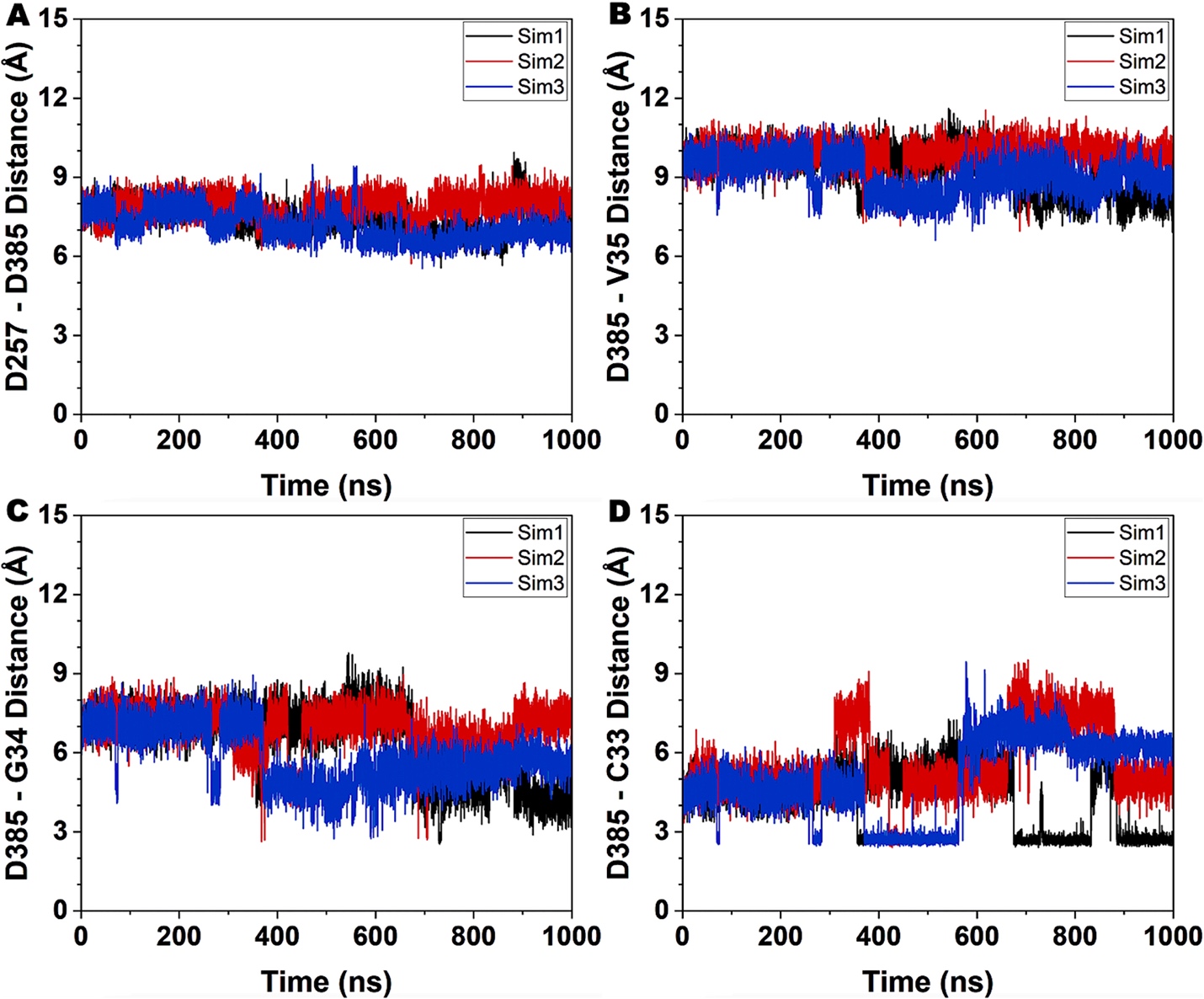

**Figure S5. The “Active-33” low-energy conformational state of model L36F Notch-bound γ-secretase compared to the “Active-34” state of model WT Notch-bound γ-secretase. (A)** The “Active-33” conformation of model L36F Notch-bound PS1 compared to the “Active-34” conformation of model WT Notch-bound PS1. **(B)** Active site of the model L36F Notch-bound PS1 in the “Active-33” low-energy conformation. A water molecule was recruited by water-bridged hydrogen bonds between the PS1 catalytic aspartates D257 and D385. **(C)** Conformation of the Notch substrate in the “Active-33” conformation of model L36F Notch-bound PS1. **(D)** Conformation of TM6a in the “Active-33” conformation of model L36F Notch-bound PS1.

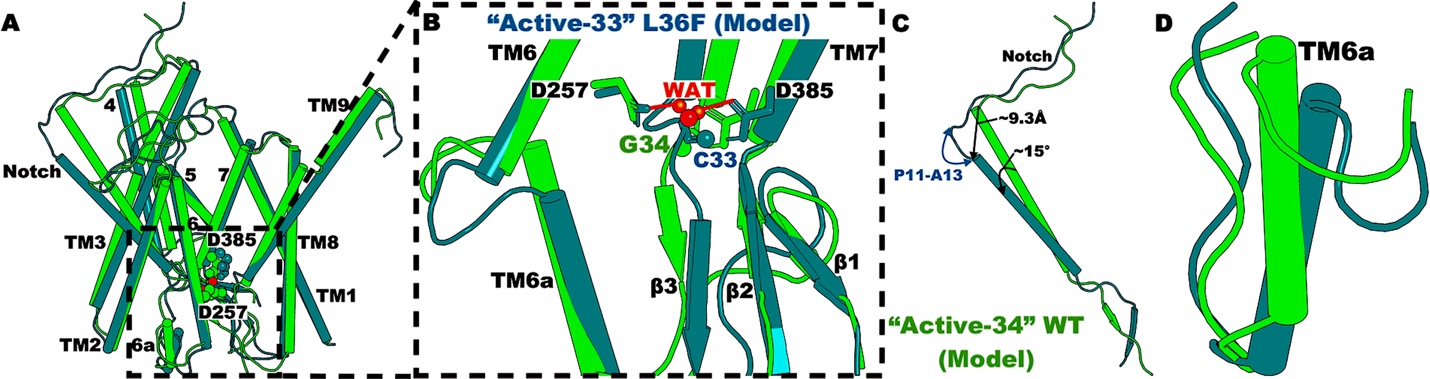

**Figure S6. The “E1” and “M1” low-energy conformational states compared to the “Active-34” state of WT Notch γ-secretase. (A)** The “E1” conformation of cryoEM WT Notch-bound PS1 compared to the “Active-34” conformation of model WT Notch-bound PS1. **(B)** Active site of the cryoEM WT Notch-bound PS1 in the “E1” low-energy conformation. The distance between PS1 residues D257 and D385 is ~6.5 Å, and the distance between PS1 residue D385 and Notch residue G34 is ~4.9 Å. **(C)** Conformation of the Notch substrate in the “E1” conformation of cryoEM WT Notch-bound PS1. **(D)** Conformation of TM6a in the “E1” conformation of cryoEM WT Notch-bound PS1. **(E)** The “M1” conformation of model WT Notch-bound PS1 compared to the “Active-34” conformation of model WT Notch-bound PS1. **(F)** Active site of the model WT Notch-bound PS1 in the “M1” low-energy conformation. The distance between PS1 residues D257 and D385 is ~6.5 Å, and the distance between PS1 residue D385 and Notch residue G34 is ~5.0 Å. **(G)** Conformation of the Notch substrate in the “M1” conformation of model WT Notch-bound PS1. **(H)** Conformation of TM6a in the “M1” conformation of model WT Notch-bound PS1.

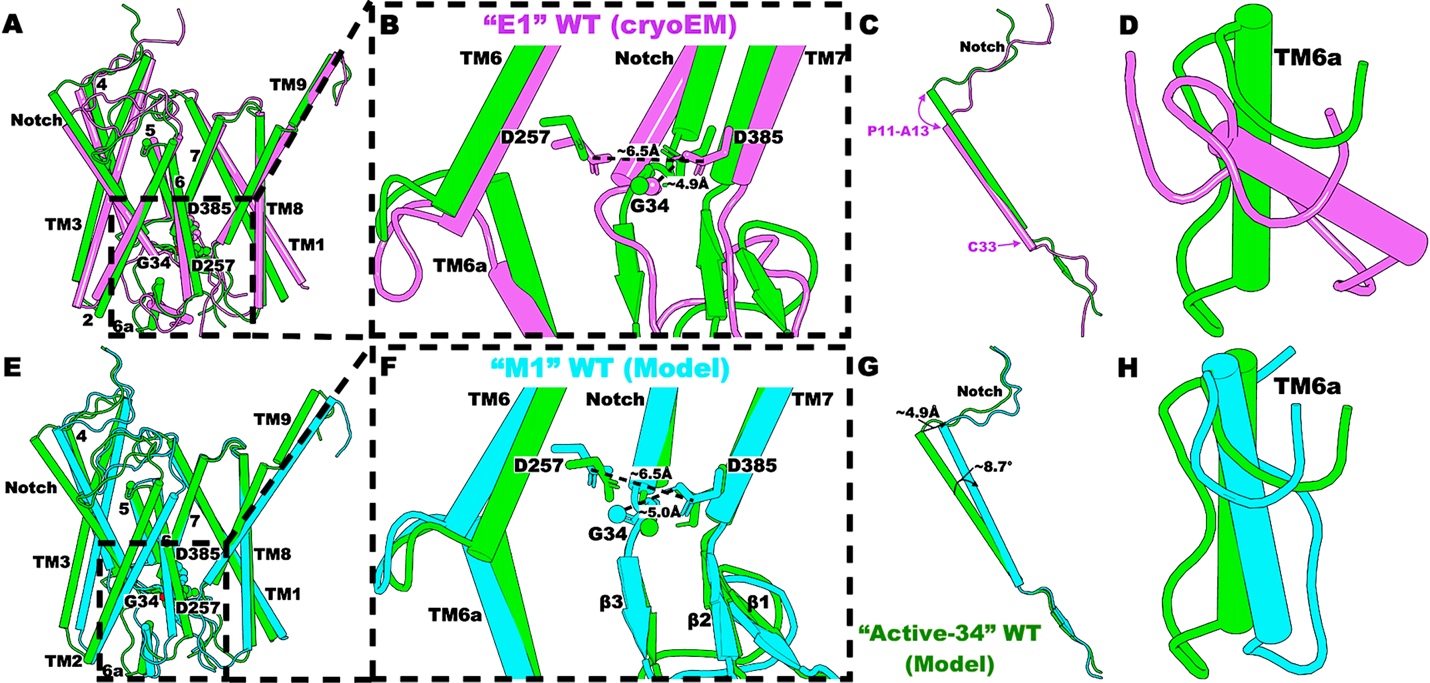

**Figure S7. The “M2” and “M3” low-energy conformational states compared to the “Active-33” state of model L36F Notch γ-secretase. (A)** The “M2” conformation of model L36F Notch-bound PS1 compared to the “Active-33” conformation. **(B)** Active site of the model L36F Notch-bound PS1 in the “M2” low-energy conformation. The distance between PS1 residues D257 and D385 is ~6.5 Å, and the distance between PS1 residue D385 and Notch residue C33 is ~6.1 Å. **(C)** Conformation of the Notch substrate in the “M2” conformation of model L36F Notch-bound PS1. **(D)** Conformation of TM6a in the “M2” conformation of model L36F Notch-bound PS1. **(E)** The “M3” conformation of model L36F Notch-bound PS1 compared to the “Active-33” conformation. **(F)** Active site of the model L36F Notch-bound PS1 in the “M3” low-energy conformation. The distance between PS1 residues D257 and D385 is ~7.2 Å, and the distance between PS1 residue D385 and Notch residue C33 is ~4.8 Å. **(G)** Conformation of the Notch substrate in the “M3” conformation of model L36F Notch-bound PS1. **(H)** Conformation of TM6a in the “M3” conformation of model L36F Notch-bound PS1.

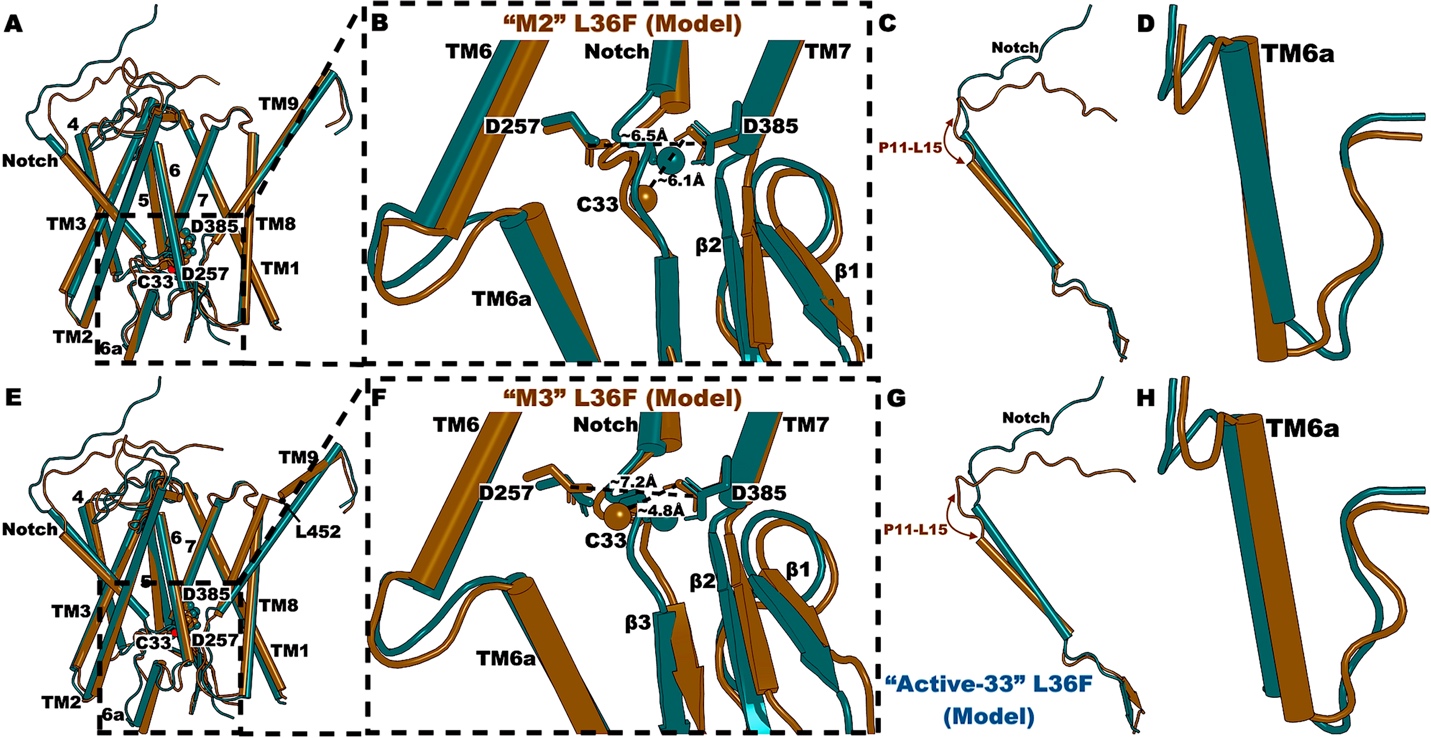

**Figure S8. Time-dependent secondary structures of model Notch bound to γ-secretase calculated from the GaMD simulations.** Time courses of the Notch secondary structures in the model WT **(A-C)** and L36F **(D-F)** Notch-bound γ-secretase calculated from GaMD simulations.

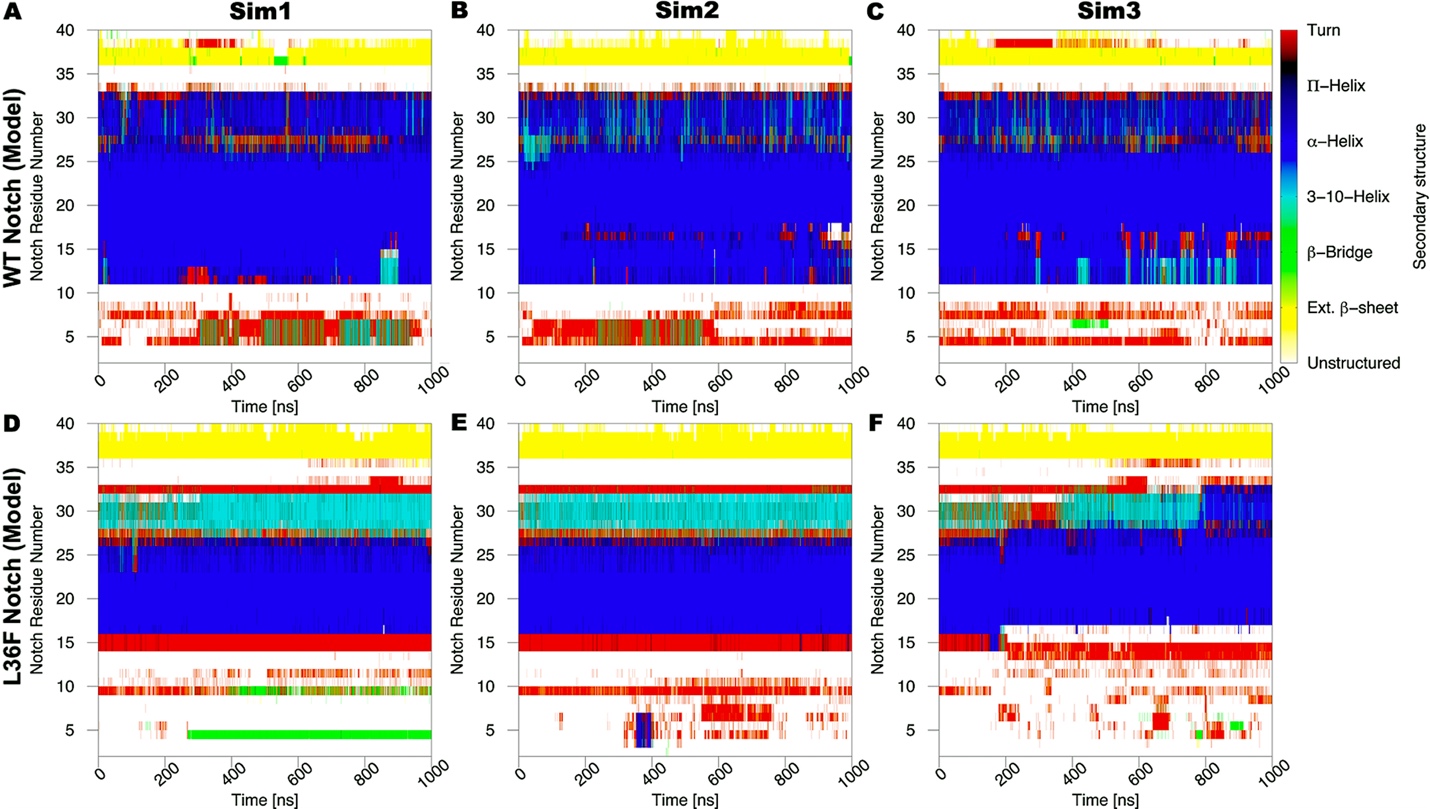

**Figure S9. Time-dependent secondary structures of cryoEM Notch bound to γ-secretase calculated from the GaMD simulations.** Time courses of the Notch secondary structures in the cryoEM WT (PDB: 6IDF) **(A-C)** and L36F **(D-F)** Notch-bound γ-secretase calculated from GaMD simulations.

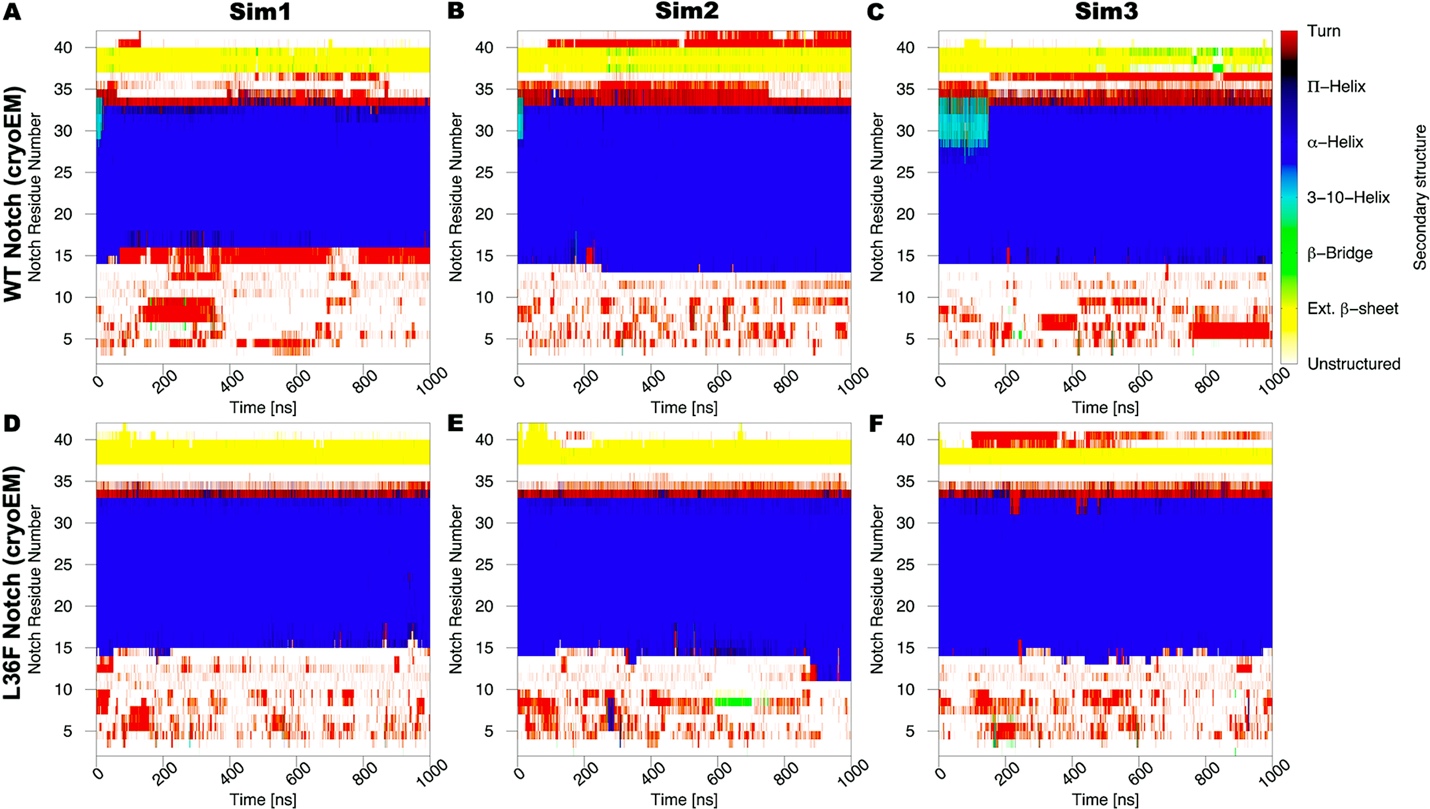

**Table S1.** Summary of GaMD simulations performed on the cryoEM and model Notch-bound γ-secretase complexes, including WT and L36F Notch-bound γ-secretase.

| **System** | **Method** | **Simulation Length** | **∆V (kcal/mol)** |
| --- | --- | --- | --- |
| cryoEM WT Notch | GaMD_Dual | 3 x 1000 ns | 14.68 ± 4.48 |
| cryoEM L36F Notch | GaMD_Dual | 3 x 1000 ns | 13.99 ± 4.38 |
| Model WT Notch | GaMD_Dual | 3 x 1000 ns | 14.04 ± 4.36 |
| Model L36F Notch | GaMD_Dual | 3 x 1000 ns | 14.94 ± 4.45 |
